# Tubulin hyperacetylation drives HMGB1 nuclear exit via the ROS-PARP1 axis leading to rotenone-induced G2/M Arrest

**DOI:** 10.1101/2025.03.25.645152

**Authors:** Sourav Dutta, Semanti Chakraborty, Ayushi Ghosh, Priyadarshini Halder, Shubhra Majumder, Ratnadip Paul, Somsubhra Nath, Piyali Mukherjee

**Affiliations:** Institute of Health Sciences, Presidency University Canal Bank Rd, DG Block, Action Area 1D, New Town, Kolkata-700156, West Bengal, India

**Keywords:** HMGB1, rotenone, Tubulin acetylation, mitochondrial ROS, Glycirrhizic acid, PJ34, G2/M arrest

## Abstract

Rotenone, a lipophilic pesticide, is strongly linked to dopaminergic neuronal loss in Parkinson’s disease (PD), primarily through mitochondrial complex I inhibition. While rotenone induces G2/M arrest in dividing cells, the underlying molecular events remain unclear. We identify HMGB1 as a key player in this process. HMGB1, known for its roles in genomic integrity and inflammation, exits the nucleus during rotenone-induced G2/M arrest, whereas its nuclear retention protects against mitotic DNA damage. We further reveal that tubulin hyperacetylation precedes HMGB1 nuclear release. Notably, reducing tubulin acetyltransferase lowers mitochondrial ROS (mtROS), preventing HMGB1 nuclear exit and mitotic DNA damage. Additionally, inhibiting PARP1 hyperactivation with PJ34 blocks HMGB1 exit and prevents G2/M arrest. These findings suggest that ROS-induced DNA damage elevates PARP1 activity, promoting HMGB1 nuclear exit and interfering with DNA repair. This tubulin acetylation/mtROS/HMGB1 axis may underlie rotenone-induced neurotoxicity, contributing to dopaminergic neuron vulnerability in PD and offering potential neuroprotective targets.

## Introduction

High mobility group box 1 (HMGB1) is a highly conserved non-histone DNA binding protein with a wide array of cellular functions that varies according to its subcellular localization (1, 2). It is the most studied member of the HMGB subfamily, contains two HMG-box domains (Box A and B) that bind DNA and bend it to facilitate interactions with other proteins and an acidic tail (3, 4). HMGB1 plays a crucial role in gene transcription, DNA repair and chromatin remodelling (5, 6). While primarily nuclear, HMGB1 can translocate to the cytoplasm and subsequently to the extracellular space acting as a damage associated molecular pattern (DAMP) (7, 8). Post-translational modifications (PTMs) play critical roles in regulating HMGB1’s sub-cellular localization and function in response to multiple cellular stress signals. For example, acetylation of lysine residues within its nuclear localization sequences (NLS) disrupts its interaction with nuclear import machinery, preventing its re-entry into the nucleus and promoting its retention in the cytoplasm (9, 10). In addition, PARP1 mediated poly-ADP ribosylation (PARylation) promotes HMGB1 cytoplasmic translocation in macrophages (11, 12). These modifications are often triggered by cellular stressors such as DNA damage, oxidative stress, or inflammatory signals. However, the implications of nuclear exit of HMGB1 under different pathological conditions triggered by non-inflammatory stimuli remains poorly understood.

Neurodegenerative diseases like Parkinson’s disease (PD) are marked by a convergence of neuroinflammation and cell cycle dysregulation, implicating a complex interplay of cellular and molecular processes that drives neuronal dysfunction (13, 14). Few studies have shown increase in HMGB1 levels both in PD patient serum samples as well as in experimental PD models such as exposure to pesticides like rotenone (15, 16). However, the role of HMGB1 in these neurodegenerative diseases is still emerging and far from understood. Rotenone, a mitochondrial complex I inhibitor, is widely used as an experimental model to study PD-related pathologies due to its ability to induce mitochondrial dysfunction, excessive reactive oxygen species (ROS) generation, and subsequent oxidative stress (17, 18). Our recent study has shown that rotenone exposure in neuronal cells leads to increased DNA damage and hyperactivation of PARP1 resulting in loss of cellular NAD^+^ levels (19).

It is hypothesized that these metabolic and oxidative perturbations may be the key drivers of DNA damage and genomic instability associated with rotenone exposure, which in turn may activate cell cycle checkpoints and induce cell cycle arrest. In line with this hypothesis, in malignant cells, rotenone induces cell cycle arrest predominantly at the G2/M phase (20, 21), which also signifies an intriguing phenomenon in the context of neuropathology. For example, dopaminergic neurons (DNs) are maintained in G2 state for prolonged periods that can lead to endoreduplication in PD, where cells replicate their DNA without division, resulting in polyploidy and genomic instability (22, 23). Mature DNs in the substantia nigra (SNc) also show aberrant cell cycle reactivation in PD. Postmortem brain analyses of DN of PD patients reveal increased phosphorylation of retinoblastoma protein (pRb), a key regulator of G1-S phase transition. (24). These events may further contribute to neuronal dysfunction and vulnerability, which are hallmarks of neurodegeneration in PD. However, to date, the mechanism underlying rotenone-induced G2/M arrest and its relationship with HMGB1 nuclear exit remains largely unexplored.

ROS plays a crucial role in regulating cytoskeletal dynamics and the DNA damage response (DDR). Elevated ROS levels, often resulting from mitochondrial dysfunction induced by environmental toxins like rotenone, can lead to tubulin hyperacetylation (25). This hyperacetylation stabilizes microtubules, enhancing intracellular transport and nuclear-cytoplasmic communication also facilitating the efficient transport of key DNA repair proteins such as p53, BRCA1, and ATM to sites of damage (26). Studies on various epithelial cells suggest that rotenone alters microtubule dynamics by directly binding to tubulin in breast and cervical cancer cells, while rotenone-induced tubulin hyperacetylation leads to the disruption of autophagic flux in a retinal pigment epithelial line (27, 28). Acetylation of α-tubulin at lysine 40 residue (K40) is catalyzed by the acetyltransferase αTAT1/Mec17, while the process is reversed by two enzymes, the NAD-independent deacetylase HDAC6 and the NAD-dependent deacetylase SIRT2. Together, these three enzymes regulate intracellular α-tubulin K40 acetylation levels (29, 30, 31).

There are limited reports suggesting the involvement of HMGB1 in cell cycle regulation and senescence, with evidence indicating its dual role in different cellular contexts. In various cells like IMR90 (human fetal lung fibroblast), MEF (mouse embryonic fibroblast), and HMEC (human mammary epithelial cells), both overexpression and depletion of HMGB1 were found to block the cell cycle and induce a senescence phenotype (32, 33). In contrast, recent studies indicate that inhibition of HMGB1 release blocks senescence in cultured astrocytes and the hTau mouse brain (34). Interestingly, knockdown of HMGB1 using siRNA results in decreased cellular proliferation, suggesting that the loss of nuclear HMGB1 may directly impact cell cycle progression (35). Additionally, HMGB1 plays an important role in base excision repair (BER) and recruits to the bulky DNA lesions induced by chemotherapeutic agents (36).

Here we investigated the role of HMGB1 focusing on two key aspects: the mechanisms driving the nuclear exit of HMGB1 following rotenone treatment and the potential involvement of HMGB1 in rotenone-induced G2/M cell cycle arrest, which may link oxidative stress to cell cycle dysregulation. We show that rotenone treatment leads to enhanced PARylation and acetylation of HMGB1, facilitating its nuclear exit, a critical event tightly associated with rotenone-induced G2/M arrest. Remarkably, inhibition of HMGB1 nuclear release effectively mitigates rotenone-induced G2/M arrest in SH-SY5Y cells, highlighting its central role in this pathway. Furthermore, we identify tubulin hyperacetylation as an upstream event preceding HMGB1’s nuclear exit via its modulation of mitochondrial ROS and subsequent DNA damage, orchestrating a novel and intricate mitochondria-to-nucleus crosstalk. This interplay of cellular events pinpoints HMGB1 as a pivotal regulator of rotenone-induced cell cycle arrest, mitotic catastrophe, and subsequent cell death which may have implications in HMGB1 mediated signalling events during disease pathology.

## Results

### Rotenone triggers time-dependent HMGB1 nuclear exit coupled with increased PARylation

HMGB1 is predominantly localized within the nucleus and acts as a DNA chaperone regulating chromatin stability and gene transcription (5, 6). While HMGB1 is known to translocate out of the nucleus in response to various cellular insults, the implications of its nuclear exit remain unclear. Notably a study has shown that the mitochondrial complex I inhibitor rotenone promotes HMGB1 nuclear exit (37) but the precise regulation along with its functional significance has not been explored in detail. Our previous study highlighted the significance of temporal regulation in cellular events following rotenone treatment (19). Therefore, we first investigated whether HMGB1 nuclear exit is also temporally regulated in response to rotenone exposure. Immunofluorescence analysis revealed that treatment of the neuronal SH-SY5Y cells with 5 μM rotenone induced time-dependent HMGB1 nuclear exit (**Fig. 1A and B**) starting at around 6 h post-treatment. Western blot analysis showed similar time-dependent cytoplasmic enrichment of HMGB1 following rotenone treatment (**Fig. 1C**) with a marked decrease in nuclear/cytoplasmic ratio of HMGB1 confirming the immunofluorescence data. HMGB1’s nuclear exit is regulated by several post-translational modifications like oxidation, PARylation and acetylation (9, 11). Since our previous study demonstrated that rotenone treatment of SH-SY5Y cells resulted in PARP1 hyperactivation leading to loss of NAD^+^ and cellular energy deficits (19), we sought to determine if this hyperactivated PARP1 results in the post-translation modification of HMGB1 via PARylation and contribute towards its nuclear exit. To investigate this, we performed a pull-down assay using an anti-HMGB1 antibody, followed by western blot analysis with an anti-PAR antibody. Our data revealed increased PARylation of HMGB1 at 24 h post-rotenone treatment (**Fig. 1D**). To validate this, we conducted a reverse pull-down assay using an anti-PAR antibody, followed by western blot analysis with an anti-HMGB1 antibody. A distinct HMGB1 band was observed exclusively in rotenone-treated cells, confirming its PARylation (**Fig. 1D**). To understand the implication of this PARylation on HMGB1 nuclear exit, we pre-treated the cells with the PARP1 inhibitor PJ34 and checked its localization both by western blot and immunofluorescence analysis. Our results reveal that PJ34 prevents rotenone-induced HMGB1 nuclear exit (**Fig. 1E and F**) indicating that PARP1 hyperactivation and HMGB1 PARylation are linked events. Next, we investigated whether HMGB1 physically interacts with PARP1, prior to its release from the chromosome. Immunofluorescence analysis revealed a notable co-localization of HMGB1 with PARP1 at 4 h post-rotenone treatment (**Fig. S1A**), suggesting a potential interaction. To confirm this, we performed co-immunoprecipitation assays using antibodies against HMGB1 and PARP1, which further validated their association following rotenone treatment (**Fig. S1B**). These findings indicate that HMGB1 interacts with PARP1 in a temporally regulated manner leading to its PARylation that potentially influencing its nuclear dynamics and subsequent exit.

**Fig. 1.**
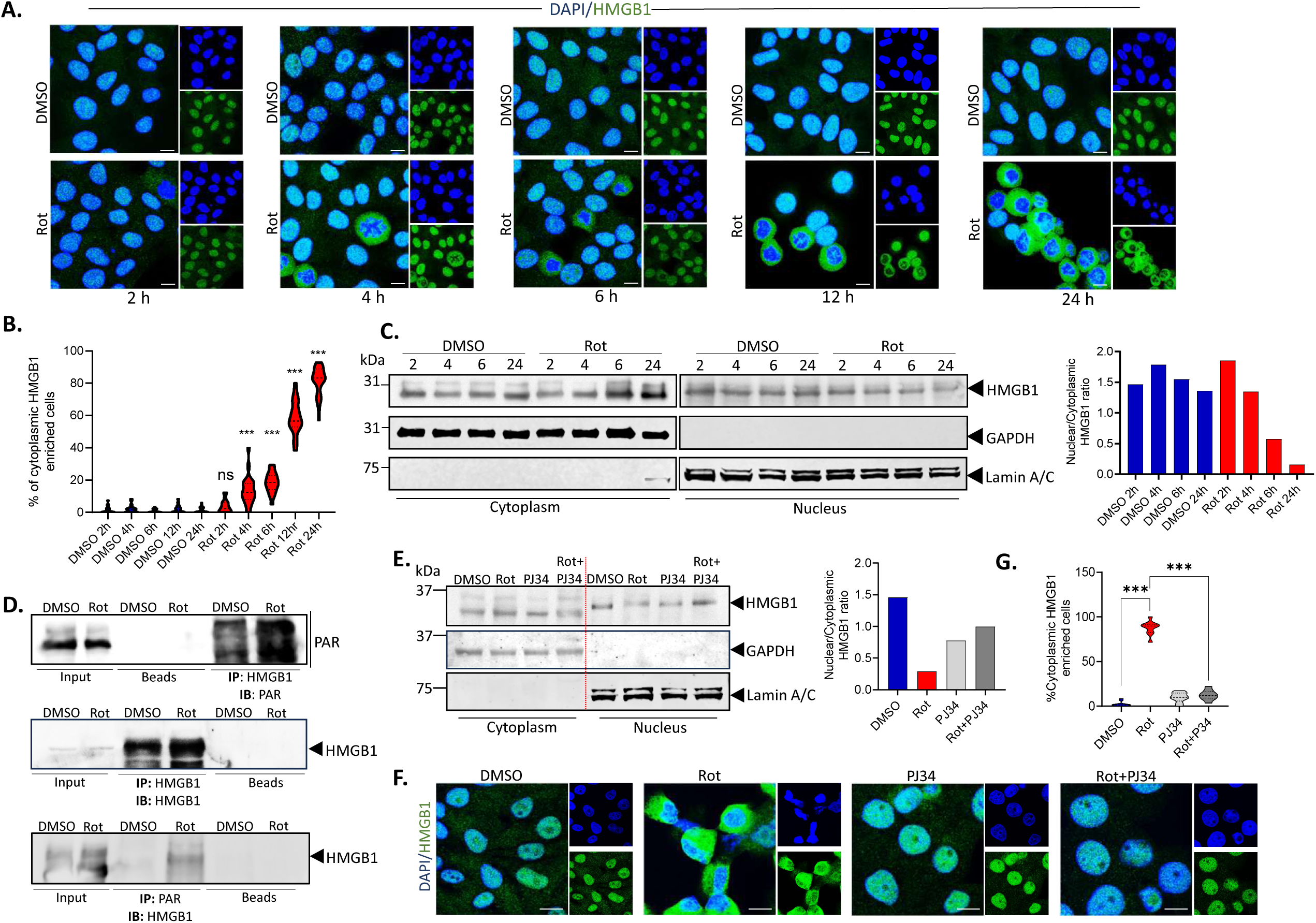
Rotenone triggers time-dependent HMGB1 nuclear exit coupled with increased PARylation. **(A)** Representative confocal images of time-dependent HMGB1 (green) localization in SH-SY5Y cells treated with rotenone (5µM) or DMSO control. DAPI (blue) indicates nuclear staining (scale bar=10 µm). **(B)** Quantification of cells with cytoplasmic enrichment of HMGB1 shown in (A). Nucleus was defined by area of DAPI. Cytosolic HMGB1 was calculated by subtracting nuclear region of interest (ROI) from the area of green channel. n=100 cells for each quantification. **(C)** Immunoblot analysis of HMGB1 protein levels in the nuclear and cytosolic fraction of cells treated with rotenone for 2, 4, 6,12 and 24 h and compared with DMSO treated controls. GAPDH was used as cytosolic loading control and lamin A/C was used as a nuclear loading control. **(D)** Co-immunoprecipitation analysis was performed to assess the PARylation status of HMGB1 in whole-cell extracts following 24 h of rotenone treatment. Cells were immunoprecipitated with an anti-HMGB1 antibody, and PAR (upper panel) and HMGB1 (middle panel) levels were analyzed. Additionally, cells were immunoprecipitated with an anti-PAR antibody, and HMGB1 levels (lower panel) were examined. **(E)** Immunoblot analysis of HMGB1 protein levels in the nuclear and cytosolic fraction of cells treated with rotenone for 24 h in the presence or absence of the PARP inhibitor PJ34. GAPDH was used as cytosolic loading control and lamin A/C was used as a nuclear loading control. **(F)** Representative confocal images of HMGB1 (green) and DAPI (blue) in SH-SY5Y cells treated with rotenone or DMSO control in the presence of PJ34. DAPI (blue) indicates nuclear staining (scale bar=10 µm). **(G)** Quantification of cells with cytoplasmic enrichment of HMGB1. n=100 cells for each quantification. Data are presented as mean ± SEM. Statistical significance was determined by one-way ANOVA with Tukey’s multiple comparison test (***P < 0.001; n=3 biological replicates).

### PARylation primes HMGB1 for acetylation following rotenone treatment that is associated with decreased SIRT1 activity

PARylation and acetylation are the key regulators of HMGB1 nuclear-cytoplasmic trafficking, facilitating its export from the nucleus (9–11). It has been suggested that PARylation weakens HMGB1’s chromatin binding and primes it for acetylation, which promotes its nuclear exit, while SIRT1 maintains HMGB1 in its deacetylated form (12) (**Fig. 2A**). We have previously observed that increased PARP1 activity leads to a depletion of NAD^+^ levels (19), potentially limiting its availability for other NAD+-dependent enzymes such as SIRT1. We hypothesized that PARP1 and SIRT1 may play opposing role in HMGB1 cellular dynamics and hence we sought to determine SIRT1 activity following rotenone treatment. We observed that the reduction in NAD^+^ levels correlates with decreased SIRT1 activity at 24 h post-rotenone treatment (**Fig. 2B**), an effect that is prevented in the presence of the PARP1 inhibitor PJ34. Consistently, we observed increased acetylation of HMGB1 in rotenone-treated cells, suggesting a potential link between reduced SIRT1 activity and enhanced HMGB1 acetylation (**Fig. 2C**). Given that SIRT1 is a key NAD^+^-dependent deacetylase, we next investigated whether rotenone-induced SIRT1 inhibition influences HMGB1 nuclear exit. To examine the time-dependent relationship between SIRT1 and HMGB1 nuclear dynamics, we assessed their localization at both early (4 h) and late (12 h) time points following rotenone treatment (**Fig. 2D and E**). Immunofluorescence analysis revealed that at 4 h post-rotenone treatment, HMGB1 and SIRT1 remained predominantly co-localized in the nucleus similar to DMSO treated controls (**Fig. 2E**). However, by 12 h HMGB1 showed increased cytoplasmic enrichment (as indicated by arrows), particularly in cells with low SIRT1 levels whereas cells with higher SIRT1 levels retained HMGB1 within the nucleus (**Fig. 2E and F**). The importance of SIRT1 in the regulation of HMGB1 nuclear exit was further strengthened by the fact that pre-incubation of cells with the SIRT1 inhibitor EX-527 along with PJ34 enhanced the cytoplasmic enrichment of HMGB1 (**Fig. 2G**) and failed to prevent rotenone-induced cell death that was achieved in the presence of prior incubation with PJ34 alone (**Fig. 2H**). The inability of PJ34 to retain HMGB1 in nucleus in the presence of a SIRT1 inhibitor points towards a critical role of functional SIRT1 in maintaining HMGB1 in a deacetylated state, thereby preventing its nuclear exit. Together, these findings reveal a novel reciprocal relationship between PARP1 and SIRT1 following rotenone treatment, wherein increased PARP1 activity depletes NAD^+^ levels, impairing SIRT1 function and leading to enhanced PARylation and acetylation of HMGB1. This post-translational modification cascade may facilitate HMGB1’s detachment from chromatin promoting its nuclear exit and contributing to rotenone-induced cellular stress.

**Fig. 2.**
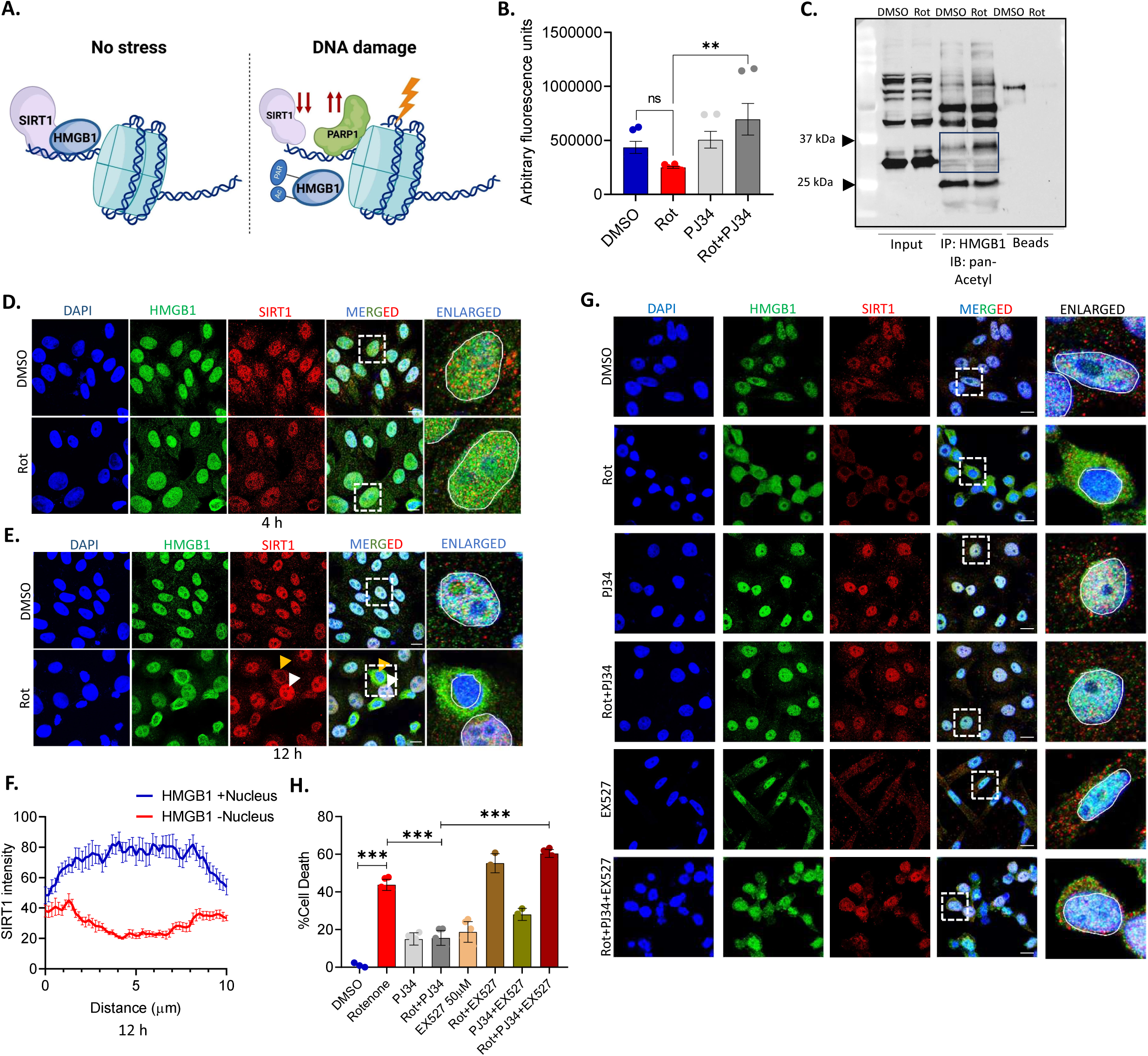
PARylation primes HMGB1 for acetylation following rotenone treatment that is associated with decreased SIRT1 activity. **(A)** Graphical representation of post-translational modification induced by PARP1 that modulates SIRT1-HMGB1 interaction during ongoing DNA damage. **(B)** Determination of SIRT1 activity by fluorometric assay in SH-SY5Y cells following rotenone treatment in the presence or absence of PJ34 and compared with DMSO-treated controls. Results are expressed in arbitrary fluorescence units. **(C)** Immunoprecipitation analysis with anti-HMGB1 antibody to check the acetylation status of HMGB1 in the whole cell extracts of cells following rotenone treatment for 12 h and probing with pan-acetyl antibody. **(D-E)** Representative confocal images of cells stained with SIRT1 (red), HMGB1 (green) and DAPI (blue) following 4 (D) or 24 (E) h rotenone treatment. (scale bar=10 µm). **(F)** Quantification of HMGB1-SIRT1 localization at 24 h post rotenone treatment. (n=100 cells for each quantification). **(G)** Representative confocal images of cells stained with SIRT1 (red), HMGB1 (green) and DAPI (blue) following 24 h rotenone treatment in the presence or absence of PJ34 or a combination of PJ34 and the SIRT1 inhibitor EX527. (scale bar=10 µm). **(H)** MTT assay of cells treated with rotenone for 24 h in the presence or absence of PJ34 or a combination of PJ34 and EX527. Data are presented as mean ± SEM. Statistical significance was determined by one-way ANOVA with Tukey’s multiple comparison test (**P < 0.01; ***P < 0.001; n=3 biological replicates).

### Inhibiting HMGB1 nuclear export by glycirrhizic acid prevents rotenone-induced G2/M cell cycle arrest

Our previous findings established that rotenone-induced PARP1 activation and impaired SIRT1 activity leads to increased PARylation and acetylation of HMGB1 and its subsequent nuclear export. Interestingly, a detailed analysis of our immunofluorescence images revealed that HMGB1 predominantly localizes to the cytoplasm in cells exhibiting intense nuclear staining suggesting condensed chromosome (**Fig. 1A**), which are seen in early mitosis, when the nuclear envelope start breaking down. Given that rotenone has been reported to induce G2/M arrest (21), we next explored whether this cell cycle perturbation correlates with HMGB1 nuclear exit. Aberrant activation of the cell cycle in post-mitotic neurons is an emerging feature of several neurodegenerative diseases, including PD, where cell cycle re-entry is associated with neurodegeneration (38). Since HMGB1 release from the nucleus is a well-known driver of cellular stress responses, we hypothesized that its nuclear export may not simply be a downstream consequence of rotenone-induced mitotic arrest but could actively contribute to this process. To test this hypothesis, we treated SH-SY5Y cells with rotenone for 24 h and FACS analysis of these cells revealed that almost 80% of these cells remain arrested in the G2/M phase of the cell cycle (**Fig. S2A**). We observed a marked increase in the mitotic cyclin, Cyclin B1, accompanied by a significant rise in the levels of Securin, a key regulator of the metaphase-to-anaphase transition (**Fig. S2B**). The accumulation of Cyclin B1, along with securin following rotenone treatment, suggests a disruption in normal mitotic progression (39, 40). Under normal conditions, as mitosis progresses, Cyclin B1 is degraded by the anaphase-promoting complex/cyclosome (APC/C) via ubiquitination, leading to MPF inactivation and facilitating mitotic exit (41). However, the rotenone-induced accumulation of Cyclin B1 strongly implies potential cell cycle arrest that could contribute to its enhanced accumulation. To further confirm this, we co-stained the cells with phospho-histone H3 (H3S10), a marker for mitotic cells, and found that rotenone-induced G2/M arrest begins as early as 6 h post-treatment **(Fig. S2C-G)**, with a significant percentage of cells (60%) becoming arrested at the prophase stage (as determined by the ring pattern of pH3S10 staining) by 12 h post treatment (**Fig. S2F-H**). Interestingly, the initiation of G2/M arrest coincided with the release of HMGB1 into the cytoplasm (**Fig. S2C-G)**, prompting us to further investigate a potential correlation between these two phenomena.

Glycyrrhizic acid (GA), a triterpenoid saponin, is known to prevent the nucleocytoplasmic translocation of HMGB1 (42) and has been shown to exert neuroprotective function in the context of neuroinflammation where HMGB1 predominantly acts as a DAMP (43). To understand the implication of the enhanced nuclear exit and cytoplasmic enrichment of HMGB1 in rotenone treated cells, we pre-incubated the cells with GA followed by rotenone treatment to prevent the nuclear exit of HMGB1. Immunofluorescence analysis revealed that GA pre-treatment of cells effectively prevents the nuclear exit of HMGB1 following rotenone treatment (**Fig. 3A and B**). Interestingly, GA pre-treatment also reduced the number of cells arrested at the G2/M phase of the cell cycle in response to rotenone treatment (**Fig. 3A and D**). This observation indicates that HMGB1 nuclear retention may play a role in preventing rotenone-induced cell cycle dysregulation. We confirmed our immunofluorescence data by performing western blot analysis of these cells and observed nuclear enrichment of HMGB1 in these cells (**Fig. 3C**). Conforming to these observations, we also observed a marked reduction in the accumulation of Cyclin B1 as well as securin levels in cells pre-treated with GA followed by rotenone treatment (**Fig. 3E**). To further confirm whether the cells are arrested at the G2/M boundary we checked the status of the mitotic checkpoint protein BubR1 which plays an important role in maintaining high fidelity during chromosome segregation (44). Our results showed that BubR1 levels are higher in rotenone treated cells but we did not see much change in its level in cells pre-incubated with GA and treated with rotenone (**Fig. 3F**) and was comparable to DMSO controls. Taken together, these data suggest that blocking the nuclear to cytoplasmic translocation of HMGB1 by GA has a significant impact on rotenone-induced G2/M arrest as depicted in **Fig. 3G**.

**Fig. 3.**
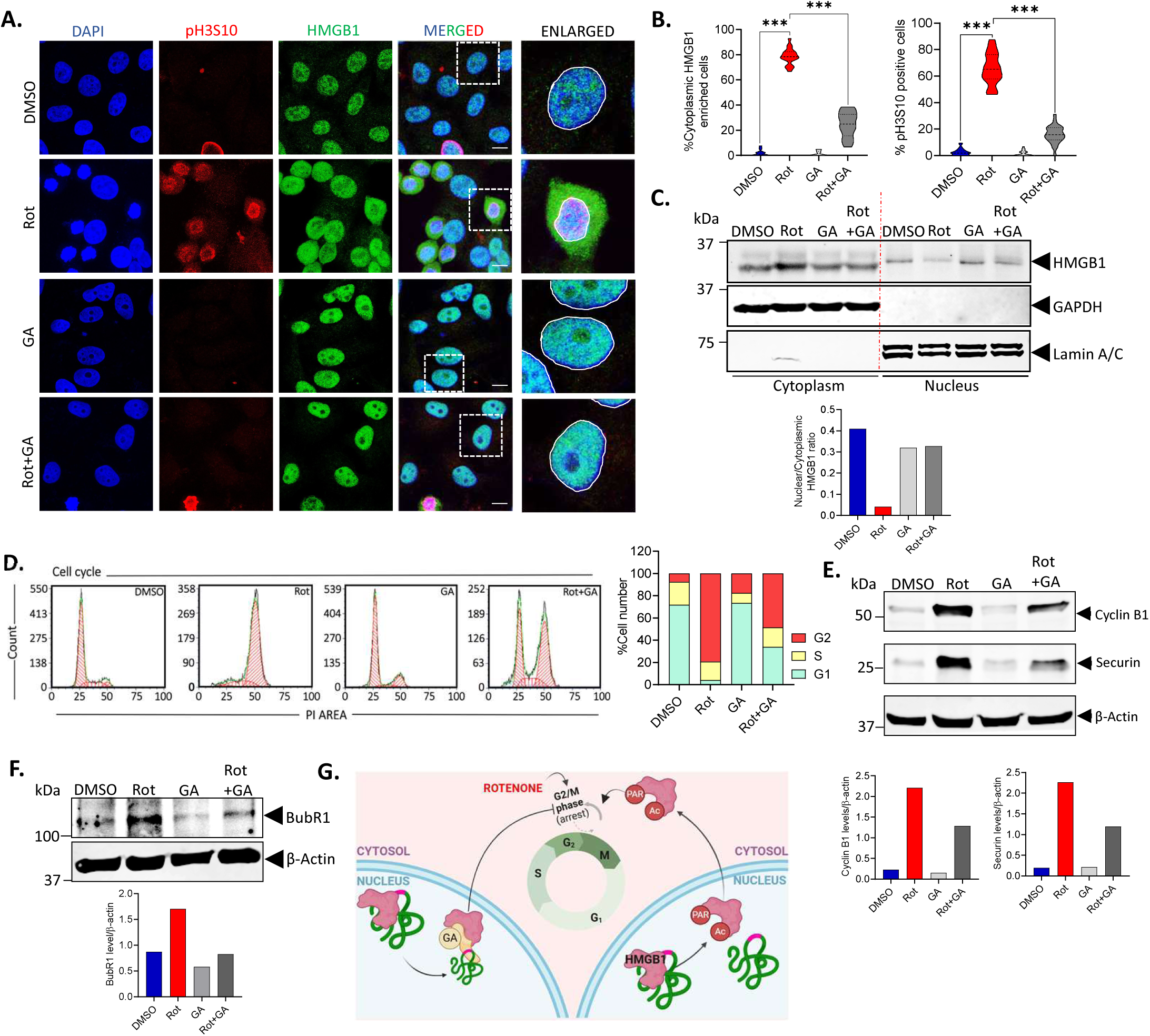
HMGB1 inhibitor glycyrrhizic acid prevents rotenone-induced G2/M arrest. **(A)** Representative confocal images of cells stained with pH3S10 (red), HMGB1 (green) and DAPI (blue) following rotenone treatment in the presence or absence of the HMGB1 inhibitor glycyrrhizic acid (GA). (scale bar=10 µm). **(B)** Quantification of cells with cytoplasmic enriched HMGB1 (left panel) and pH3S10 positive cells indicating cells arrested at G2/M (right panel) for each treatment condition. (n=100 cells for each quantification). **(C)** Immunoblot analysis of HMGB1 protein levels in the nuclear and cytosolic fraction of cells treated with rotenone for 24 h in the presence or absence of GA. GAPDH was used as cytosolic loading control and lamin A/C was used as a nuclear loading control. **(D)** Representative flow cytometry histograms depicting DNA content in cells stained with propidium iodide (PI). Distinct peaks correspond to cells in the G0/G1 phase (first peak), S phase (intermediate region), and G2/M phase (second peak). SH-SY5Y cells were treated with Rot, GA or pre-incubated with GA before Rot treatment for 24 h and the data is compared with DMSO control. The quantification of cells in each phase of the cell cycle are shown in the right panel **(E-F)** Immunoblot analysis of Cyclin B1, Securin (E) and BubR1 (F) protein levels in the whole-cell extracts of SH-SY5Y cells 24 h post-rotenone treatment in the presence or absence of GA. β-Actin was used as a loading control. **(G)** Schematic representation showing association of rotenone-induced G2/M arrest with HMGB1 nuclear exit and the effect of GA on the process. Data are presented as mean ± SEM. Statistical significance was determined by one-way ANOVA with Tukey’s multiple comparison test (***P < 0.001; n=3 biological replicates).

### HMGB1 hypoacetylation mutants prevents its nuclear exit and subsequent rotenone-induced G2/M arrest

We previously demonstrated that rotenone induces HMGB1 acetylation, and earlier studies have shown that HMGB1 acetylation is critical for its cytoplasmic translocation. This prompted us to investigate whether blocking HMGB1 acetylation could prevent its nuclear exit and whether this impacts rotenone-induced G2/M arrest. HMGB1 contains two nuclear localization signals (NLS): **NLS1**, located in the A box (amino acids 28–44), and **NLS2**, downstream of B box (amino acids 179–185). NLS1 includes four conserved lysine residues, and NLS2 has five conserved lysine residues, both susceptible to acetylation and are known to influence HMGB1 nuclear export. To explore this, we generated several HMGB1 hypoacetylation mutants, as illustrated in **Fig. 4A**. The lysine residues in NLS1 (K28–30) and NLS2 (K182–185) were mutated to alanine or arginine to create single or double NLS mutants. SH-SY5Y cells were transfected with these mutants for 24 h, followed by rotenone treatment for another 24 h and stained with anti-FLAG and anti-pH3S10 antibodies. Immunofluorescence analysis revealed that in the hypoacetylation mutants, HMGB1 was retained in the nucleus in conformation with a previous study (45), and this nuclear localization was unaffected by rotenone treatment (**Fig. 4B and C**). Interestingly, this nuclear retention of HMGB1 prevented rotenone-induced G2/M arrest specifically in these cells, as indicated by the absence of pH3S10 staining in the transfected cell population. To further validate our findings, we generated a nuclear-enriched version of HMGB1 by deleting its putative nuclear export signal (NES), which spans amino acid residues 118–132. Consistent with our hypothesis, the majority of NES-deleted mutants were retained in the nucleus, and these cells did not exhibit signs of G2/M arrest (**Fig. 4B**). However, a subset of cells still displayed extracellular HMGB1, which correlated with increased pH3S10 staining, a hallmark of mitotic entry. This suggests that additional, yet unidentified, NES sequences in HMGB1 may contribute to its nuclear exit under rotenone-induced stress conditions and requires further investigation. In summary, these novel findings reinforce our earlier observations with GA, suggesting that HMGB1 nuclear exit is closely associated with rotenone-induced G2/M arrest that could be prevented by enriching nuclear HMGB1.

**Fig. 4.**
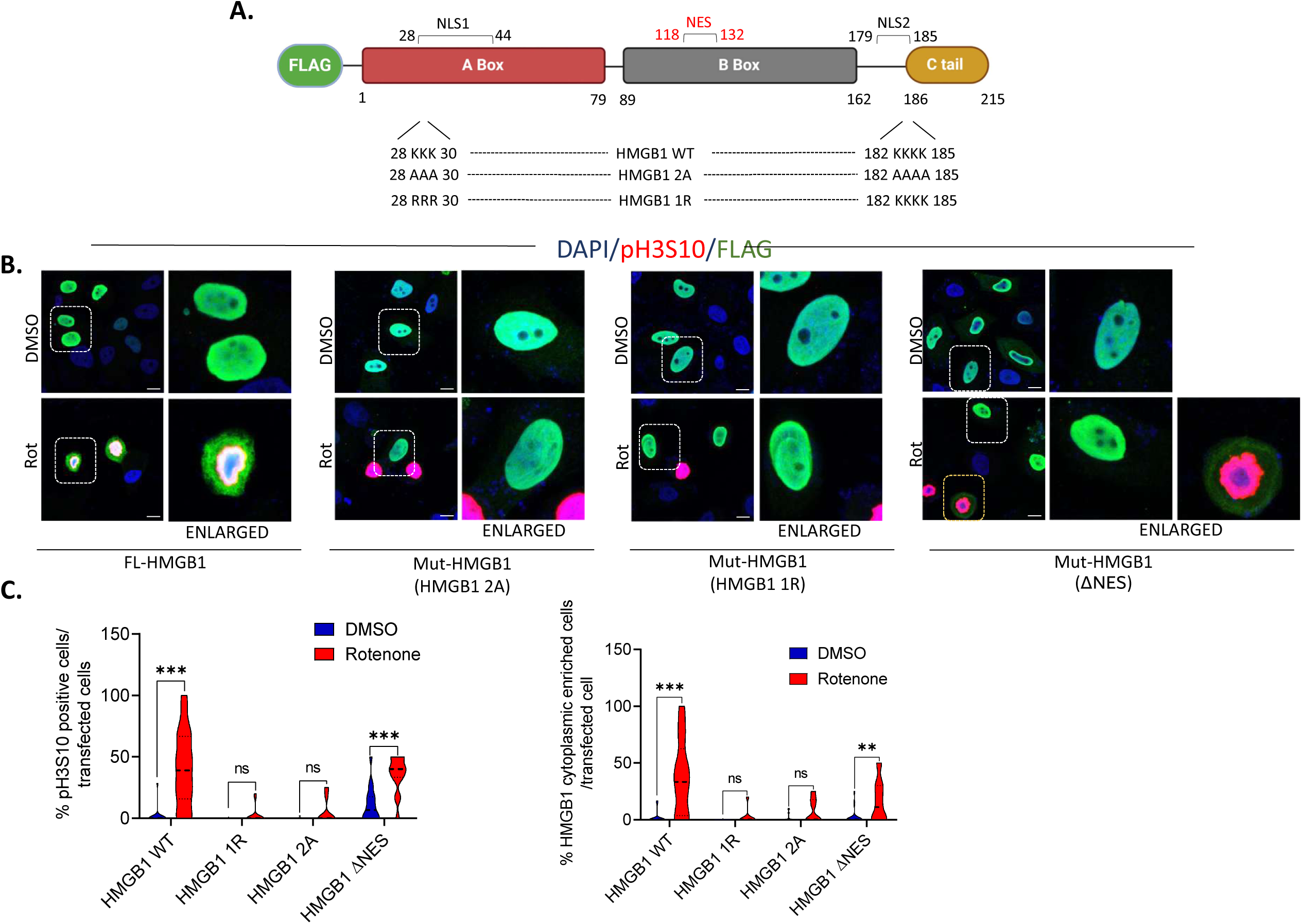
HMGB1 hypoacetylation mutants prevents its nuclear exit and subsequent rotenone-induced G2/M arrest. **(A)** Schematic illustration of sites mutated in HMGB1 for this study. **(B)** Representative confocal images of cells transfected with FLAG-tagged wild-type HMGB1 or its mutants as shown in (A) and immunostained with FLAG (green), pH3S10 (red) and DAPI (blue) following rotenone treatment for 24 h. (scale bar=10 µm). **(C-D)** Quantification of pH3S10 positive cells per transfected cell (C) and cells with cytoplasmic enriched HMGB1 per transfected cell. (n=50 transfected cells for each quantification). Data are presented as mean ± SEM. Statistical significance was determined by two-way ANOVA with Tukey’s multiple comparison test (***P < 0.001).

### Rotenone-induced tubulin hyperacetylation triggers mtROS-induced DNA damage that precedes HMGB1 nuclear release

We next sought to understand how nuclear enrichment of HMGB1 following rotenone treatment prevents G2/M cell cycle arrest. A key area of interest is the disruption of microtubule dynamics, a known cause of mitotic arrest, though no study till date have linked this mechanism to HMGB1 cellular dynamics. Since rotenone has been shown to bind to tubulin, we analyzed the status of microtubules in rotenone treated SH-SY5Y cells. Our findings reveal that rotenone induces robust tubulin hyperacetylation in these cells as early as 4 h post-treatment, with levels increasing over time (**Fig. 5A**). To further dissect whether this hyperacetylated tubulin impacts the nucleocytoplasmic translocation of HMGB1, we employed αTAT1 (alpha tubulin acetyl transferase 1) knock down cells for further analysis. Rotenone treatment in these αTAT1 knockdown (KD) cells for 24 h, markedly reduced tubulin acetylation (**Fig. S3A-B**). Interestingly, αTAT1 KD cells showed enriched nuclear HMGB1 despite rotenone exposure (**Fig. 5B, Lower panel and 5C, Upper panel**), unlike control cells (**Fig. 5B, Upper panel and 5C, Upper panel**), suggesting a role for hyperacetylated tubulin in rotenone-induced G2/M arrest. Supporting this, rotenone failed to induce G2/M arrest in αTAT1 KD cells (**Fig. 5B, Lower panel**), further linking HMGB1 nuclear retention and rotenone-induced cell cycle arrest. To further dissect this relationship, we pre-treated cells with the HMGB1 inhibitor GA, and found that while GA blocked HMGB1 nuclear exit and G2/M arrest as previously demonstrated (**Fig. 4**), it did not affect tubulin hyperacetylation (**Fig. 5D**). To further confirm this, we treated cells with nocodazole, a well-established microtubule-depolymerizing agent that disrupts microtubule dynamics and arrests cells in mitosis. As expected, nocodazole treatment did not induce tubulin hyperacetylation at 24 h post-treatment. Importantly, GA pre-treatment failed to prevent G2/M arrest in nocodazole-treated cells (**Fig. 5D and E**), indicating that HMGB1-mediated G2/M arrest is specific to rotenone exposure. In conclusion, these findings suggest that tubulin hyperacetylation occurs upstream of HMGB1 release but is insufficient to drive G2/M arrest in the presence of nuclear-retained HMGB1. Taken together, these results highlight a complex interplay between microtubule acetylation, HMGB1 sub-cellular localization, and rotenone-induced G2/M arrest.

**Fig. 5.**
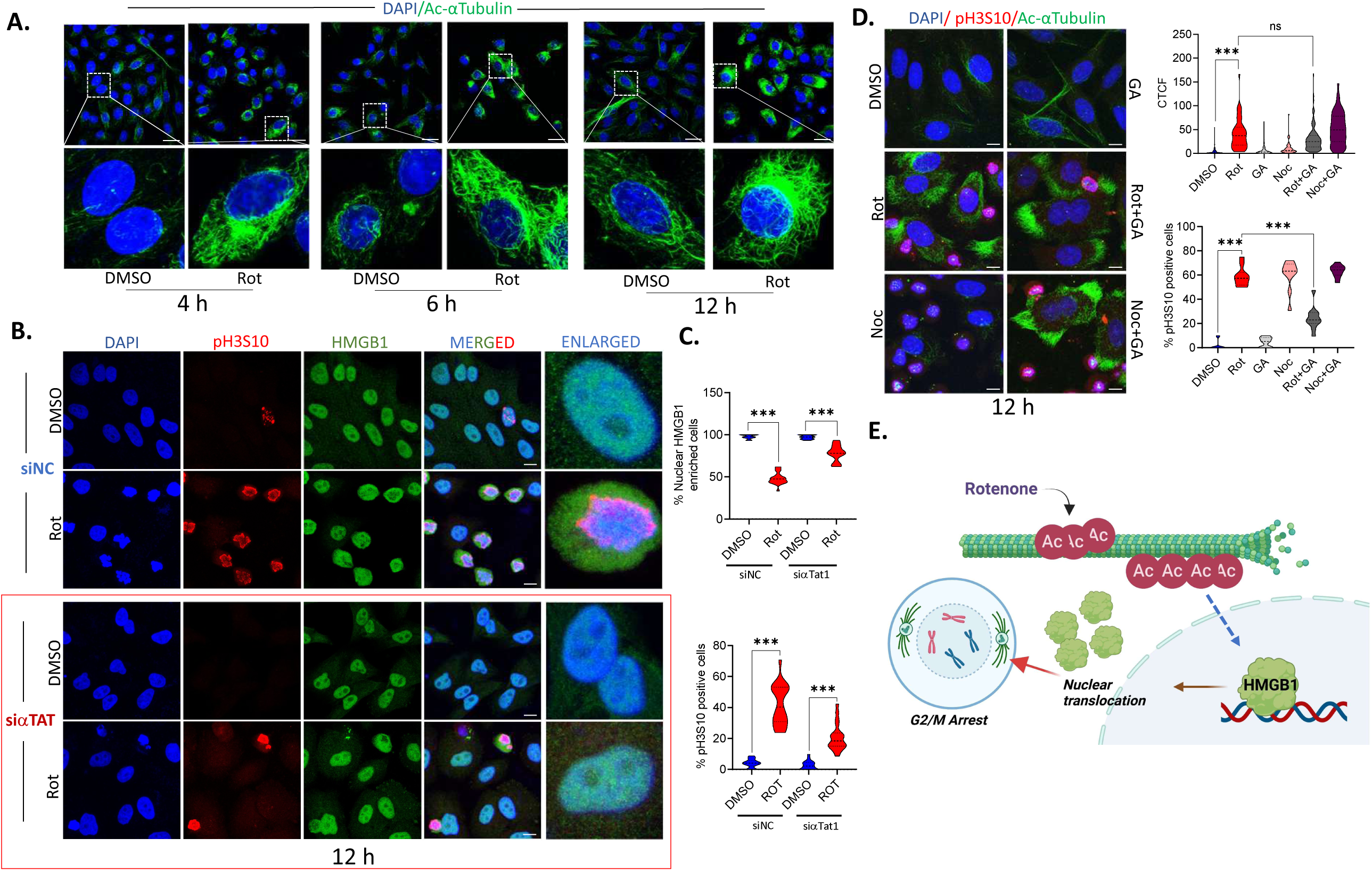
Rotenone-induced tubulin hyperacetylation precedes HMGB1 nuclear release. **(A)** Representative confocal images of cells stained with Ac-αTubulin (green) and DAPI (blue) following rotenone treatment for 4, 6 and 12 h (scale bar=25 µm). **(B)** Representative confocal images in cells stained with pH3S10 (red), HMGB1 (green) and DAPI (blue) in cells transfected with control siRNA (siNC) or siαTAT1. (scale bar=10 µm). **(C)** Quantification of percentage of cells with nuclear enriched HMGB1 (upper panel) and cells arrested at G2/M as depicted by pH3S10 positive cells (lower panel). (n=50 transfected cells for each quantification). **(D)** Representative confocal images depicting the effect of GA on the acetylation of α-tubulin (green) induced by rotenone alongwith with pH3S10 (red) and DAPI (blue). (Scale bar= 10 µm). Corrected total cell fluorescence (CTCF) values of α-tubulin intensity of control vs rotenone or nocodazole treated cells in the presence or absence of GA (right upper panel). Quantification of pH3S10 positive cells (right lower panel) similarly treated. (n=100 transfected cells for each quantification). **(E)** Schematic representing the effect of tubulin hyperacetylation on HMGB1 nuclear exit and subsequent G2/M arrest induced by rotenone. Data are presented as mean ± SEM. Statistical significance was determined by one-way ANOVA with Tukey’s multiple comparison test (***P < 0.001; n=3 biological replicates).

To further investigate the mechanism by which tubulin hyperacetylation drives HMGB1 nuclear exit, we explored the source of rotenone-induced cellular stress. Rotenone, a mitochondrial complex I inhibitor, is known to induce mitochondrial dysfunction and elevate mitochondrial ROS (mtROS) levels. Previous studies have shown that tubulin acetylation plays a crucial role in maintaining microtubule stability, intracellular trafficking, and mitochondrial dynamics (46). Notably, hyperacetylation of tubulin has been linked to both protective and detrimental effects on mitochondrial function, depending on the cellular context and is an emerging area of interest. A key study demonstrated that various cellular stresses, including oxidative stress induced by hydrogen peroxide (H₂O₂), lead to microtubule hyperacetylation (25). Here we hypothesized that mitochondrial dysfunction induced by rotenone and tubulin hyperacetylation are closely linked events that could promote HMGB1 nuclear exit. To investigate this further, we assessed mtROS levels using MitoSOX staining in rotenone-treated siControl and siαTAT1 cells. Our results show that rotenone-induced tubulin hyperacetylation correlated with increased mtROS production as mtROS levels were substantially reduced in the siαTAT1 cells both at 4 and 12 h post treatment (**Fig. 6A and B**). Since in our previous study we observed that pre-treatment of cells with the antioxidant NAC could not reverse rotenone-induced cell death (9) or motor deficits in rotenone-exposed flies (47), we re-examined its impact on HMGB1 release and found no effect on rotenone-induced nuclear exit of HMGB1 (**Fig. S4 A & B**). However, the earlier observation of reduction of mtROS in the absence of tubulin hyperacetylation intrigued us. A deeper investigation into the published literature showed that although NAC could reverse mitochondrial disruptions induced by lower doses of rotenone, it could not reduce mtROS level (48). This prompted us to use another inhibitor of mitochondrial superoxide generation Diphenyleneiodonium (DPI) (49) and as depicted in **Fig. 6C**, pre-treatment of SH-SY5Y cells with DPI at a dose of 2.5 μM, substantially reduced mtROS alongwith rotenone-induced HMGB1 nuclear exit (**Fig. 6D and E, Left panel**). Interestingly, consistent with our previous findings, DPI-treated cells also exhibited reduced pH3S10 staining, particularly in cells with enriched nuclear HMGB1 (**Fig. 6D and E, Right panel**).

**Fig. 6.**
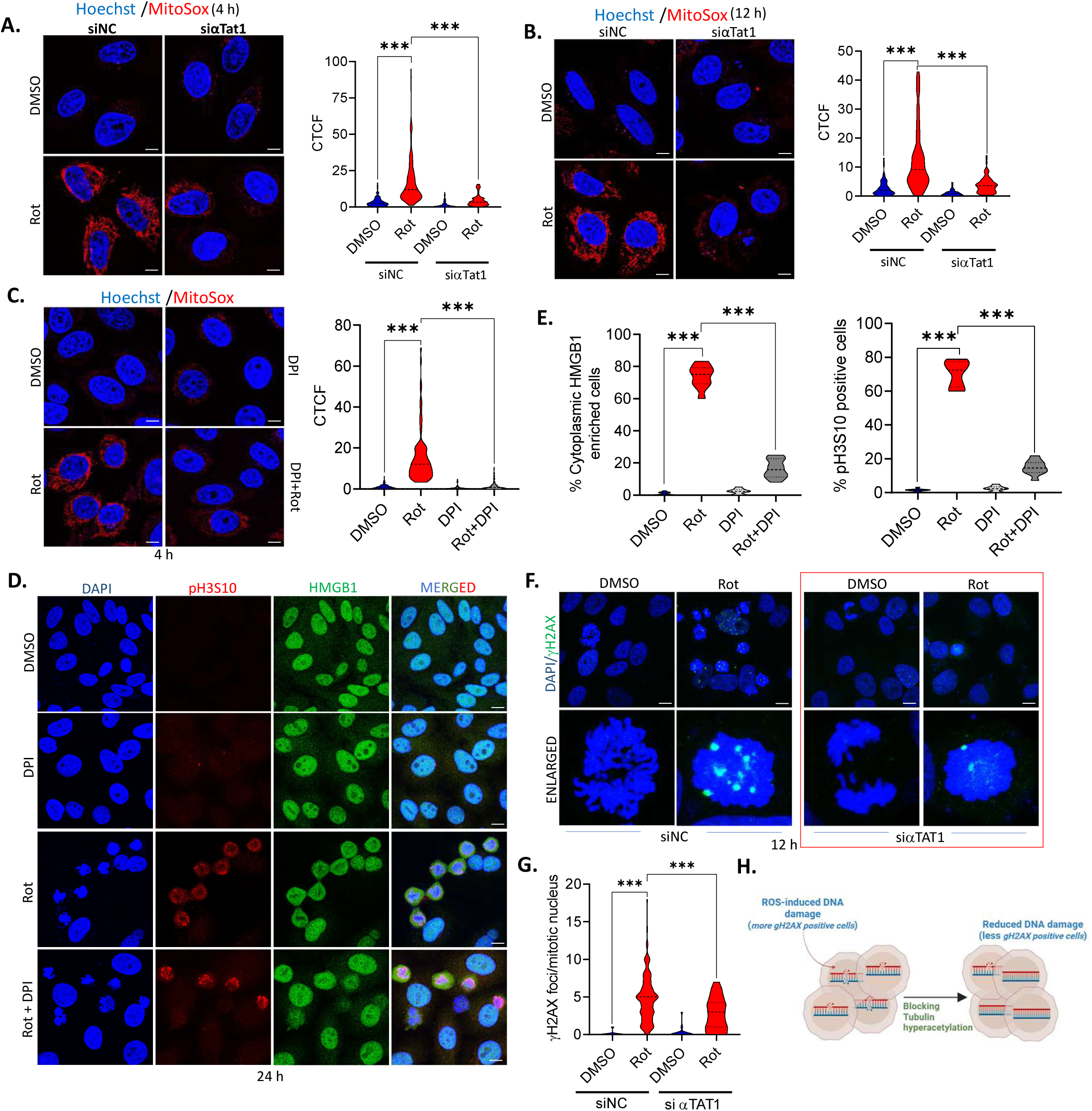
Rotenone-induced tubulin hyperacetylation triggers mtROS-induced DNA damage. **(A-B)** Visualization of mitochondrial ROS (mtROS) production by live staining with the mitochondria-specific ROS indicator MitoSox red (red) and the nuclear stain Hoechst 33342 (blue) at 4 h (A) and 12 h (B) post-rotenone treatment in cells transfected with control siRNA (siNC) or siαTAT1. The CTCF values of MitoSox intensity are also shown. (scale bar=10 µm). **(C)** Visualization of mtROS production by staining with MitoSox red (red) and the nuclear stain Hoechst 33342 (blue) at 12 h post-rotenone treatment in the presence of absence of the NAD(P)H inhibitor DPI (2.5 µM). (scale bar= 10 µm). **(D)** Representative confocal images of pH3S10 (red), HMGB1 (green) and DAPI (blue) following rotenone treatment in the presence or absence of DPI. (n=100 cells for each quantification; scale bar=10 µm). **(E)** Quantification of cytoplasmic enriched HMGB1 (left panel) and pH3S10 positive cells indicating cells arrested at G2/M (right panel) and for each treatment condition for 100 cells. **(F)** Representative confocal images of cells stained with phospho-γH2AX (green) and DAPI (blue) following rotenone treatment in siNC and siα-TAT1 cells. (Scale bar=10 µm). **(G)** Quantification of γH2AX positive cells in rotenone treated cells transfected with siControl and α-TAT1 knockdown. (n=50 cells for each transfection and treatment). **(H)** Schematic representation showing the effect of mtROS in inducing ds-breaks in nuclear DNA following rotenone treatment that is alleviated by blocking tubulin acetylation. Data are presented as mean ± SEM. Statistical significance was determined by one-way ANOVA with Tukey’s multiple comparison test (***P < 0.001; n=3 biological replicates).

Although our data demonstrated a correlation between tubulin hyperacetylation and increased mtROS levels following rotenone treatment, which in turn appeared to be involved in HMGB1 nuclear exit, the underlying mechanistic details remained unclear. Our earlier study has shown that rotenone triggered early PARP1 hyperactivation that could be due to increased DNA damage associated with rotenone treatment. Further, our data convincingly demonstrated that HMGB1 undergoes PARylation that primes it for nuclear exit following rotenone treatment and could be prevented by the PARP inhibitor PJ34 (**Fig. 2**). Since we observed that decreased mtROS production was linked to HMGB1 nuclear enrichment, we further evaluated its impact on rotenone-induced DNA damage. As depicted in **Fig. 6F and G**, there was a significant decrease in DNA damage foci in rotenone-treated siαTAT1 cells compared to the siControl cells as evident by number of γH2AX positive nuclei which is a well-established marker of DNA double-strand breaks. This clearly indicates that reducing tubulin acetylation mitigates rotenone-induced mtROS production and subsequent HMGB1 nuclear exit alongwith associated DNA damage. In conclusion, our data provides a critical link wherein tubulin hyperacetylation primes mtROS-mediated DNA damage and subsequent PARylation-driven HMGB1 nuclear release, thereby contributing to rotenone-induced cellular stress responses and subsequent G2/M arrest as demonstrated in **Fig. 6H**.

### Rotenone induces mitotic catastrophe that could be correlated to an impaired DNA damage response in the absence of nuclear HMGB1

When cells detect DNA damage, they typically arrest at the G2/M boundary to facilitate repair (50). However, prolonged arrest at this checkpoint or during early mitosis can lead to mitotic catastrophe, ultimately resulting in cell death (51). Our analysis of pH3S10 staining indicated that rotenone-treated cells were unable to progress beyond prophase, suggesting a sustained mitotic block that could result due to ongoing DNA damage. To determine whether these cells could re-enter the cell cycle after rotenone withdrawal, we treated them with rotenone for 24 h and then released them for an additional 12 h (schematic in **Fig. 7A**). Interestingly, rotenone treated cells remained arrested at the G2/M boundary even after release, as evident by persistent pH3S10 staining (**Fig. 7B**). In contrast, cells treated with nocodazole, another mitotic blocker, successfully re-entered the cell cycle upon drug removal, showing a significant reduction in mitotic cell numbers (**Fig. 7B**). Furthermore, cells released from rotenone treatment exhibited persistent DNA damage compared to those treated with nocodazole (**Fig. 7D**), suggesting that rotenone treated cells are unable to recover from the G2/M arrest that may ultimately lead to mitotic catastrophe. This was further confirmed by western blot analysis of Cyclin B1 and Securin levels at 12 h post withdrawal of rotenone or nocodazole. Consistent with pH3S10 staining, rotenone-treated cells showed elevated levels of both Cyclin B1 and Securin even after rotenone withdrawal whereas these levels significantly dropped in the nocodazole withdrawn cells (**Fig. 7D**).

**Figure 7.**
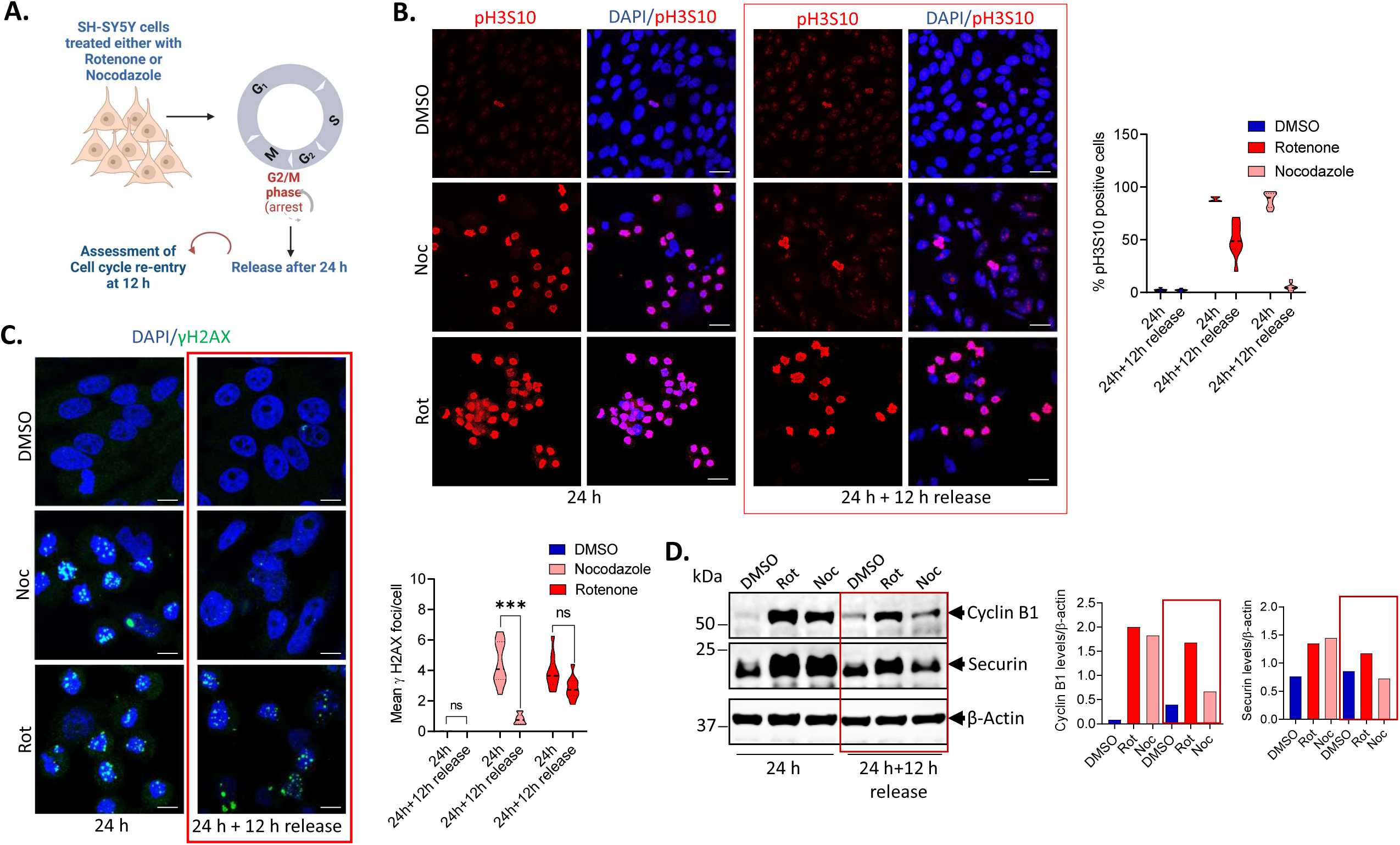
Rotenone causes persistent DNA damage that cannot be reversed following its withdrawal. **(A)** Schematic outline of experimental strategy to study mitotic release in the presence of rotenone or nocodazole **(B)** Representative confocal image of cells fixed at 12h post-release following 24h treatment with rotenone or nocodazole and stained with pH3S10 (red) and DAPI (blue). (n=100 cells for each quantification). (scale bar=25 µm). **(C)** Representative confocal images of cells stained with phospho-γH2AX (green) and DAPI (blue) fixed at 12h post-release following 24 h treatment with rotenone or nocodazole (left panel). Quantification of phospho-γH2AX positive cells (right panel) (n=100 cells for each quantification). (scale bar=10 µm). **(D)** Immunoblot analysis of Cyclin B1 and Securin protein levels in the whole-cell extracts at 12h post-release following 24 h treatment with rotenone or nocodazole. β-Actin was used as a loading control. Data are presented as mean ± SEM. Statistical significance was determined by two-way ANOVA with Tukey’s multiple comparison test (***P < 0.001; n=3 biological replicates).

Our next question was to understand why the rotenone-treated cells are unable to recover from persistent G2/M arrest and re-enter the cell cycle. We speculated that HMGB1 could regulate DNA repair genes based on its well-established role as a chromatin-binding protein involved in DNA damage sensing and repair. Previous studies have shown that HMGB1 interacts with key DNA repair proteins, facilitates chromatin remodeling, and influences the recruitment of repair factors to sites of DNA damage (52). Given that rotenone exposure leads to persistent DNA damage and mitotic arrest, we hypothesized that the loss of nuclear HMGB1 following rotenone treatment might impair the transcriptional regulation of DNA repair genes, thereby compromising the cell’s ability to recover from rotenone-induced genotoxic stress. Although HMGB1 is known to recruit DNA damage repair proteins to the site of damage, whether it modulates their expression is not known. To address this question, we re-analyzed the HMGB1 ChIP-Seq data from available resources in HUVEC cell lines (GSM2589815) and IMR90 cell line (GSM5234116) (33, 53) (**Fig. 8A**). The ChIP-Seq analysis of HMGB1 binding to the chromosome in these cell lines demonstrates its significant association with key DNA repair genes, suggesting a regulatory role in maintaining genomic integrity. Prominent HMGB1 enrichment was observed at the promoter region of **FEN1** (61,792,800 to 61,797,260 on chromosome 11), a critical player in base excision repair and replication fork maintenance. In IMR90 cells, binding peaks were detected at the **XPA** locus, spanning 97,654,398 to 97,697,630 on chromosome 9, a gene essential for nucleotide excision repair (NER), suggesting that HMGB1 might be involved in chromatin remodeling for the NER machinery recruitment. HMGB1 was also enriched at the **GADD45A** locus (67,685,100 to 67,688,334 on chromosome 1) in HUVEC cells, implicating HMGB1 in stress-responsive pathways involving DNA repair, cell cycle arrest, and apoptosis. Binding was further detected at the **KLF4** locus (107,484,852 to 107,491,360 on chromosome 9), a gene involved in genomic stability and transcriptional regulation of repair pathways, and at **RAD23B** (107,283,160 to 107,332,194 on chromosome 9), which is integral to NER. These data reveal that HMGB1 preferentially binds to the promoter and regulatory regions of critical DNA repair genes, emphasizing its role in orchestrating cellular responses to DNA damage and maintaining genomic stability under stress conditions especially those of the NER machinery. In alignment with these findings, we observed reduced expression of these genes at 24 h (**Fig. 8B**) post-rotenone treatment implicating there could be transcriptional loss in these crucial genes that could lead to an overall impairment of the DNA repair machinery. Interestingly, at 4 h post-treatment when there was substantial HMGB1 enrichment, the expression of GADD44A and KLF4 was markedly increased but subsequently declined at 24 h, coinciding with HMGB1 nuclear exit. The importance of nuclear HMGB1 in DNA damage repair process was further strengthened by the observation that pre-treatment with GA, which prevents HMGB1 nuclear exit, led to a reduction in mitotic DNA damage in rotenone-treated cells (**Fig. 8C**). In conclusion, these findings suggests that nuclear retention of HMGB1 may either protect against rotenone-induced mitotic DNA damage or facilitate DNA repair (**Fig. 8D**), warranting further investigation.

**Fig. 8.**
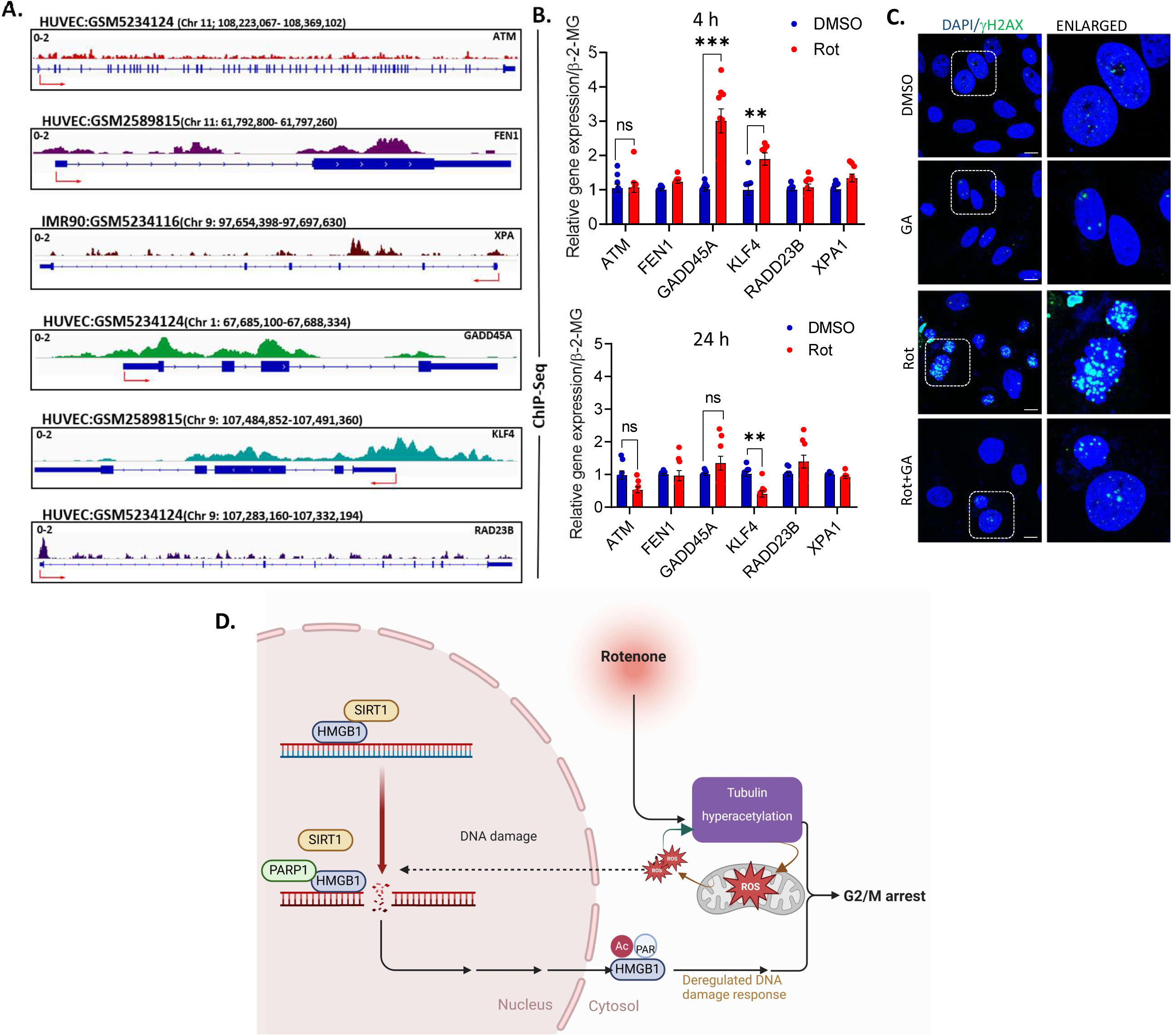
Loss of nuclear HMGB1 affects the transcription of DNA damage response genes that may lead to rotenone-induced mitotic catastrophe. **(A)** Representative IGV snapshots displaying ChIP-seq binding peaks of HMGB1, reanalysed from ChIP-Atlas database, on selected DNA Damage Response (DDR) genes ATM, FEN1, XPA, GADD45A, KLF4, RAD23B. Peaks indicate strong enrichment of HMGB1 binding, suggesting potential regulatory roles at these loci. Chromosomal position and transcription start site (red arrow) depicted in the image. **(B)** qRT-PCR analysis of the relative gene expression of DDR genes at 4 h (upper panel) or 24 h (lower panel) post-rotenone treatment with respect to DMSO-treated controls. β2 microglobulin (β2MG) was used as an internal control. **(C)** Representative confocal images of cells stained with phospho-γH2AX (green) and DAPI (blue) following rotenone treatment in the presence or absence of GA. (scale bar=10 µm). **(D)** Graphical representation of the study. Data are presented as mean ± SEM. Statistical significance was determined by unpaired t-test with Sidak multiple comparison (**p < 0.01; ***p < 0.001; n= 3 biological replicates).

## Discussion

HMGB1 is a highly conserved non-histone chromatin binding protein that has multifactorial role within the cells. As a nuclear protein, HMGB1 ensures genomic stability, facilitates DNA repair, and influences gene transcription. However, under conditions of cellular stress, HMGB1 can translocate from the nucleus to the cytoplasm where it modulates autophagy or be released extracellularly, where it functions as a damage associated molecular pattern (DAMP) triggering heightened inflammatory responses (5–8). It interacts with receptors like RAGE and TLR4, triggering inflammatory responses that are meant to contain damage but can also contribute to disease pathology (54, 55). While inflammation is a necessary part of injury response, chronic or excessive activation can accelerate neurodegeneration, as seen in diseases like Alzheimer’s and Parkinson’s (56, 57). Thus, the nuclear-cytoplasmic shift of HMGB1 represents a critical molecular switch that influences cell fate decisions between repair and apoptosis under various pathological conditions.

Despite extensive studies on HMGB1’s role in inflammation, the overall consequence of its nuclear exit in the context of cellular homeostasis and stress adaptation remains poorly understood. Here we show that the mitochondrial complex I inhibitor rotenone, triggers the nuclear exit of HMGB1 in neuronal cells which are known to be the non-immune mediators of cellular responses. Since our previous study has strongly indicated that rotenone-induced cellular energy deficits are associated with neuronal stress responses and subsequent cell death (19), we sought to determine whether HMGB1 plays a distinct role in this process.

The nuclear exit of HMGB1 is tightly regulated by post-translational modifications (PTMs), including acetylation, PARylation, phosphorylation, and glycosylation. Although these modifications have been extensively studied in immune cells or under immunological triggers such as LPS (58, 59), their roles in non-immune cells, including neurons remain largely unidentified. In this study, we report for the first time that rotenone-induced PARP1 hyperactivation in neuronal SH-SY5Y cells enhances HMGB1 PARylation. This finding raises two key questions: what are the functional consequences of HMGB1 PARylation, and does it promote HMGB1 nuclear exit following rotenone treatment? The well-established connection between PARP1 hyperactivation and NAD^+^ depletion prompted us to investigate whether NAD^+^ loss influences other NAD^+^-dependent nuclear enzymes, particularly SIRT1, which is known to maintain HMGB1 in a deacetylated state, preventing its release from chromatin. As expected, rotenone treatment significantly reduced SIRT1 activity, leading to increased HMGB1 acetylation, a modification associated with its nuclear exit. Interestingly we found that the PARP1 inhibitor PJ34 effectively prevented HMGB1 release, but this effect was abolished when the SIRT1 inhibitor EX527 was added, indicating that both PARylation and acetylation are essential for HMGB1 nuclear exit under rotenone-induced stress conditions. These findings highlight a complex regulatory network governing HMGB1 nuclear-cytoplasmic trafficking, which may have significant implications for cellular responses to mitochondrial dysfunction and oxidative stress.

One of the central questions that arose from our study is the functional significance of HMGB1 nuclear exit in the context of rotenone-induced cellular stress. While previous research has established that rotenone induces G2/M arrest, the precise molecular regulators of this arrest remain poorly characterized. Since HMGB1 is known for its DNA repair activity, we hypothesized that HMGB1 nuclear retention may confer a protective advantage to proliferating cells by enabling them to bypass or recover from rotenone-induced stress and subsequent cell cycle arrest. Consistent with this, our results demonstrated that cells with enriched nuclear HMGB1 exhibited reduced G2/M arrest, suggesting that its nuclear retention mitigates rotenone-induced checkpoint activation. To validate this hypothesis, we used two complementary approaches-firstly, pharmacological inhibition using glycyrrhizic acid (GA), which prevents HMGB1 nuclear exit and secondly, we employed genetic modification through hypoacetylation mutants of HMGB1, which restrict its localization to the nucleus. In both cases, retention of HMGB1 in the nucleus correlated with reduced G2/M arrest, highlighting its crucial role in preventing rotenone-induced cell cycle arrest. It may be noted here that the association of HMGB1 with chromatin during mitosis has been debated, with previous studies reporting conflicting findings (60, 61). However, our data provide compelling evidence that HMGB1 nuclear exit may act as a key driver of the mitotic arrest observed in rotenone-treated neuronal cells.

Building on the observation of rotenone-induced G2/M arrest and the protective effects of nuclear retention of HMGB1 in the process, we investigated early events preceding HMGB1 nuclear exit. Microtubule dynamics, particularly the acetylation of lysine 40 (K40) on α-tubulin, play a critical role in cell cycle progression. This modification, regulated by the opposing actions of histone deacetylase 6 (HDAC6) and tubulin acetyltransferase (αTAT1), serves as a marker of microtubule stabilization. While rotenone is known to induce tubulin depolymerization (24), emerging evidence suggests it also promotes tubulin hyperacetylation (25), potentially influencing cell cycle regulation and stress responses. Consistent with this, we found that tubulin hyperacetylation occurs as early as 4 h after rotenone exposure, preceding HMGB1 release. This finding raises intriguing questions about the role of microtubule modifications in HMGB1 translocation and cell cycle adaptation under stress. Consistent with this, αTAT1 knockdown (KD) cells showed marked reduction in cells arrested at G2/M following rotenone treatment that is accompanied by nuclear retention of HMGB1. These cells also exhibited decreased mtROS production and reduced mitotic DNA damage. Notably, pre-treatment with GA did not prevent rotenone-induced tubulin hyperacetylation, suggesting that this modification occurs upstream of HMGB1 nuclear exit and may serve as the initiating event for the G2/M arrest. However, the mechanistic link between tubulin hyperacetylation and HMGB1-mediated cell cycle regulation remained unresolved till this point. Hyperacetylated tubulin has been shown to recruit Drp1 leading to enhanced mitochondrial fission and increased ROS production that may exacerbate oxidative DNA damage (62). Our data suggests that tubulin hyperacetylation triggers mtROS production and excessive ROS levels can sustain tubulin in a hyperacetylated state, potentially creating a feedback loop that exacerbates oxidative stress. It is evident that this persistent ROS elevation leads to DNA damage in rotenone treated cells subsequently triggering PARP1 hyperactivation as part of the cellular response to genotoxic stress. Given that PARP1 plays a crucial role in DNA damage sensing and repair, its hyperactivation result in excessive PARylation of nuclear proteins, including HMGB1. This modification could facilitate HMGB1 nuclear exit, further linking tubulin acetylation, ROS accumulation, and DNA damage response pathways.

Prolonged mitotic arrest beyond a critical threshold often leads to a phenomenon termed mitotic catastrophe, an irreversible process that often culminates in cell death and may arise due to aberrant or non-functional DNA damage machinery (63, 64). Our data indicate that cells exposed to rotenone fail to re-enter the cell cycle even after its withdrawal as compared with another G2/M blocker nocodazole. These cells showed persistent mitotic DNA damage even after rotenone withdrawal which maybe due to an impaired DNA damage repair machinery, making recovery ineffective. Moreover, HMGB1 nuclear exit, which we observed following rotenone exposure, may further impair the DNA damage response, as nuclear HMGB1 is known to regulate key DNA repair pathways.

This was confirmed by a reanalysis of HMGB1 ChIP-Seq data which revealed that HMGB1 binds to the promoters of several genes involved in the DNA damage response (DDR) under stress conditions. This finding aligns with previous studies demonstrating HMGB1’s role as a chromatin-associated protein that modulates gene expression and DNA repair processes under oxidative stress. Intriguingly, we observed a reduction in mitotic DNA damage in both siαTAT1 and glycyrrhizic acid (GA)-pre-treated cells, prompting us to investigate whether HMGB1’s nuclear binding is essential for regulating these DNA repair genes. We observed that rotenone failed to upregulate the expression of these genes suggesting that the nuclear presence of HMGB1 is crucial for initiating DNA repair responses. This also raises the intriguing possibility that nuclear loss of HMGB1 impairs the cellular ability to recover from rotenone-induced oxidative damage, potentially leading to prolonged arrest and mitotic catastrophe. Future investigation could explore whether HMGB1’s binding to DDR gene promoters facilitates chromatin remodeling or recruits additional repair factors, providing mechanistic insights into its role in oxidative stress adaptation.

In conclusion, we propose a novel connection between tubulin hyperacetylation, mitochondrial superoxide production, and HMGB1 nuclear release, ultimately driving rotenone-induced G2/M arrest. This arrest likely represents a cellular attempt to counteract DNA damage; however, substantial loss of nuclear HMGB1 appears to disrupt this process, pre-disposing cells to mitotic failure and death. Given that neurons are post-mitotic and do not undergo conventional cell cycle progression, the implications of HMGB1 nuclear release in this context remain unclear. Notably, neurons can activate aberrant cell cycle pathways under stress, a phenomenon linked to neurodegeneration (65). Our findings highlight the protective role of nuclear HMGB1 in preventing G2/M-associated stress responses, which may have therapeutic relevance in dopaminergic neuronal death triggered by rotenone. Targeting HMGB1 nuclear retention or modulating its interactions with cell cycle regulators could offer promising strategies to protect neurons from stress-induced damage and death.

## Acknowledgements

The authors thank Abhik Saha (Presidency University, Kolkata) for critical insights during the course of the study and Debanjan Mukhopadhyay (Presidency University, Kolkata) for providing reagents. The work was funded by extramural funding from Science and Engineering Research Board core research grant (SERB-CRG), Govt. of India (#CRG/2020/001325) and Department of Science and Technology and Biotechnology, Govt. of West Bengal [2291(Sanc.)/STBT-13015/7/2024-WBSCST SEC] to PM. SD is a recipient of CSIR-NET fellowship [08/155(0085)-2020-EMR-I] from Govt. of India. RP is a recipient of ICMR Senior Research Fellowship [3/1/3/JRF-2019/HRD (LS)]. PH is a recipient of UGC-NET fellowship [F.82-1/2018 (SA-III)]. Images used in the models were created in Biorender.com.

## Author contribution

PM conceived the work and SD and SC performed the experiments. AG assisted in the construction and validation of HMGB1 mutants. PH and SM assisted with the tubulin acetylation experiments and RP and SN assisted with the mitotic analysis. SD, SC and PM wrote the manuscript and prepared the figures and PM revised the entire manuscript.

## Experimental procedures

### Cell line, antibodies and chemicals

Human neuroblastoma cell line SH-SY5Y was a kind gift from Oishee Chakrabarti (Saha Institute of Nuclear Physics, Kolkata). Cells were cultured in high glucose Dulbecco’s Modified Eagle’s Medium (DMEM; #12100046; Gibco, Thermo Fisher Scientific) supplemented with 10% FBS (#10270106; Gibco, Thermo Fisher Scientific), 1% Penicilin-Streptomycin solution (#15140122; Gibco, Thermo Fisher Scientific), 100μg/ml Gentamicin solution (#A010; Himedia) at 37°C and 5% CO2.

Mouse monoclonal antibodies against HMGB1; PCRP-HMGB1-4F10 was deposited to the DSHB by Common Fund – Protein Capture Reagents Program (DSHB Hybridoma Product PCRP-HMGB1-4F10), GAPDH; DSHB-GAPDH-2G7 was deposited to the DSHB by DSHB (DSHB Hybridoma Product DSHB-GAPDH-2G7) and LAMIN A/C MANLAC2(10F8) was deposited to the DSHB by Morris, G.E. (DSHB Hybridoma Product MANLAC2(10F8)) were purchased from Developmental Studies Hybridoma Bank (DSHB), created by the NICHD and maintained at the University of Iowa, Department of Biology, Iowa City, IA 52242. Rabbit polyclonal antibodies against HMGB1 (#10829-1-AP), and pan-acetyl antibody (#66289-1-Ig) were purchased from Proteintech. Rabbit monoclonal antibody against SIRT1 (#ab32441, E104) was obtained from Abcam. For determining the level of PARylation, anti-PAR antibody (#MCA-1480) and mouse monoclonal antibody for β-actin (#MCA5775GA) was obtained from Bio-Rad. Rabbit monoclonal antibody against PARP1 (#9532) was obtained from Cell Signaling Technology Inc. (CST). Mouse monoclonal antibody against FLAG (#F1804) was obtained from Sigma-Aldrich. Rabbit polyclonal antibodies against Securin (#13445), Cyclin B11 (#4138), BUBR1 (#4116) and Phospho-Histone 3 (Ser10) (#9701) were obtained from Cell Signaling Technology Inc. (CST). H2A.XS139 (phospho Ser139) antibody (#GTX628789) was obtained from GeneTex. Mouse monoclonal antibodies for K40 acetylated α-tubulin (6-11B-1) (#sc-23950) and α-tubulin (#sc-32293) were purchased from Santa Cruz Biotechnology. Rabbit isotype IgG (#02-6102) and Mouse isotype IgG (#02-6502) for immunoprecipitation were purchased from Thermo Fisher Scientific. HRP tagged anti-mouse (#ab97023) and anti-rabbit secondary antibodies (#ab97051) were purchased from Abcam, anti-mouse DyLight800 (#5257P) was purchased from Cell Signaling Technology Inc. and Alexa Fluor 680 anti-rabbit (#A10043; Thermo Fisher Scientific) was used. For immunofluorescence staining, secondary antibodies Alexa Fluor 594 anti-rabbit (#ab150080) and Alexa Fluor 488 anti-mouse (#ab150113) from Abcam were used.

Mitochondrial complex I inhibitor rotenone (#R8875; Sigma-Aldrich), ROS scavenger N-acetyl-cysteine (#A9165, Sigma-Aldrich), Nocodazole (#M1404; Sigma-Aldrich) and HMGB1 inhibitor Glycyrrhizic acid ammonium salt from *glycyrrhiza root (licorice)* (#50531; Sigma-Aldrich) were used for this study. PARP inhibitor PJ34 (#HY-13688A; Medchemexpress) SIRT1 inhibitor EX527 (#ab141506; Abcam), was purchased from Abcam. Diphenyleneiodonium (DPI) was a kind gift from Dr. Debanjan Mukhopadhyay (Institute of Health Sciences, Presidency University, Kolkata).

### Preparation of WT-HMGB1 constructs

For HMGB1 wild-type (WT) expression construct, pCMV6-Entry mammalian expression vector (Origene #PS100001; a kind gift from Karin Peterson, Rocky Mountain Laboratories, NIAID, NIH, USA) was used. mRNA was isolated from SH-SY5Y cells by standard PureZOL (# 7326890; Bio-Rad) method and cDNA was prepared using iScript cDNA synthesis kit (Bio-Rad, #1708841) as per the manufacturers protocol. HMGB1 ORF were amplified with the following sets of primers as described in **Table S1** using Q5 high-fidelity DNA polymerase (#M0491; NEB) as per the manufacturers protocol and cloned into pCMV6-Entry mammalian expression vector by standard digestion-ligation method. Plasmids were confirmed for the ORFs by Sanger’s sequencing method.

### Site directed mutagenesis of HMGB1

HMGB1 hypoacetylation NLS mutant constructs were made using pCMV6-FLAG-HMGB1 WT plasmid as template. Substitution of 28-30 and 182-185 lysine residues with alanine or arginine and NES (118–132) deletion mutant were done by PCR with the primers listed in the **Table S1** using Q5 high-fidelity DNA polymerase. PCR products were digested with DpnI (#RS176S; NEB) followed by addition of T4 polynucleotide kinase (#M0201S; NEB) and circularized with T4 DNA ligase (Promega #M180A) sequentially. Introduction of the mutant nucleotides were confirmed by Sanger’s sequencing method.

### Plasmid transfection

Coverslips were placed in wells of 24-well cell culture plates (Greiner Bio-One) and treated with membrane matrix Geltrex (#A1569601; Thermo Fisher Scientific) for 1 h at 37° C, and 0.02 x 10^6^ cells were seeded. After 24 h, cells were transfected with 0.75 μg DNA/well using JetOptimus (Polyplus) (#101000006) transfection reagent at 1:2 (DNA: Transfection reagent) ratio according to manufacturer’s instruction. After 24 h of transfection, medium was replaced by fresh complete growth medium and the subsequent treatments were done after 24 h of transfection.

### siRNA transfection

siRNA for αTat1 was purchased from Qiagen (FLEXITUBE GG; ID:SI03124660|S0). As per the experimental requirement 0.01 x 10^6^, 0.02 x 10^6^ or 0.1 x 10^6^ cells were seeded in a 35 mm glass bottom dishes (Cellvis), 24-well plate or 6-well cell culture dishes (Greiner; Bio-One) respectively. Transfection was done with 20nM siRNA using Lipofectamine RNAiMAX (#13778075; Thermo Fisher Scientific) according to manufacturer’s instruction. After 24 h of transfection cells were replenished with fresh growth medium and treated with rotenone or DMSO.

### Nuclear fractionation of cells

2.5 x 10^6^ cells were seeded in 100 mm dishes (Nunc; Thermo Fisher Scientific). After 24 hours, cells were treated as per the experimental requirements. Cells were collected and cytoplasmic lysis was done in 300 μl of Buffer A (10 mM HEPES pH 7.9, 10 mM KCl, 1.5 mM MgCl2, 340mM sucrose, 10% glycerol) containing 0.1% Triton X-100, 1X protease inhibitor cocktail (#78429; Thermo Fisher Scientific), 1mM PMSF, 100μM Na3Vo4, 50mM NaF and incubated for 30 min on ice with periodic mixing. After cytoplasmic lysis, cells were centrifuged at 4400 rpm for 5 min and the pellet was stored, and the supernatant was further centrifuged at 14000 rpm for 10 min twice; to remove any contaminant and the final supernatant (cytoplasmic fraction) was stored for further use. The pellet acquired after cytoplasmic lysis was resuspended in 300 μl Buffer B (3 mM EDTA, 0.2 mM EGTA) with 0.1% Triton X-100, 1X protease inhibitor, 1mM PMSF, 100μM Na3Vo4, 50 mM NaF and incubated for 30 min on ice with periodic mixing and centrifuged at 4400 rpm for 5 min. The final pellet (nuclear fraction) was dissolved in 300 μl buffer A. No vortex was done at any step of the protocol.

### Immunoblot analysis

0.3 x 10^6^ cells were seeded in the wells of a 6-well cell culture plate for 24 h and treatment done for defined time points. Cells were washed in ice cold PBS after treatment and lysed in RIPA buffer (50 mM Tris-HCL pH 7.4, 150 mM NaCl, 0.5% sodium deoxycholate, 1% NP-40, 0.1% SDS, 1mM EDTA). 1X Protease inhibitor and 1mM PMSF was added to the buffer just prior to cell lysis. Cells were lysed with periodic vortexing for 30 minutes on ice. Protein concentration was estimated using BCA protein estimation kit (#2603100011730; Bangalore GeNei) and 30μg of protein/well were resolved by SDS-PAGE following which proteins were transferred to PVDF membrane (Bio-Rad). After blocking for 1 h with 5% non-fat dry milk or 3% BSA, membranes were probed with indicated primary antibodies followed by incubation with HRP or IR conjugated secondary antibodies. Bands were detected with Chemiluminescent ECL substrate (Bio-Rad) in ChemiDoc MP imaging system (12003154; Bio-Rad) or fluorescence in LI-COR^®^ Odyssey^®^ DLx Imaging System respectively. Band intensity was quantified by ImageJ software (Developed at the National Institutes of Health and the LOCI, University of Wisconsin, Madison, WI, USA) and was plotted in GraphPad Prism 8.0.2as a ratio to the housekeeping protein.

### Co-immunoprecipitation assay

2.5 x 10^6^ cells were seeded in 100 mm dishes and after 24 h (70% confluency) cells were treated as required. Cells were harvested and cell lysis was done using 400 IP lysis buffer (25 mM Tris-Cl pH 7.4, 150 mM NaCl, 1% NP-40) freshly supplemented with 1X Protease inhibitor cocktail and 1 mM PMSF. Lysate was collected by centrifugation at 14000 rpm for 3 min. 1-1.5 mg of cell lysate was precleared and stored on ice. 2 μg primary antibody for HMGB1 (DSHB), PAR and 1:200 PARP1 was incubated with protein G (#161-4023; Bio-Rad) or A magnetic beads (#161-4013; Bio-Rad) for 2 h at room temperature by rotation and beads were washed with IP wash buffer (25 mM Tris-Cl pH 7.4, 150 mM NaCl, 0.1% Tween-20) for 10 min thrice. Precleared lysate was incubated with antibody-bead complex overnight at 4°C with rotation. Next day beads were washed with IP wash buffer for 10 minutes thrice and the beads were boiled with 2X Laemmli buffer and the supernatant was collected and analyzed by SDS-gel electrophoresis.

### Immunofluorescence analysis

12 mm coverslips were coated with Geltrex (#A15696-01; Thermo Fisher Scientific) for 1 h at 37°C and washed with 1X PBS. 0.04 x 10^6^ cells were seeded on precoated coverslips for 24 h before treatment. After treatment cells were washed with 1X PBS and fixed with 4% paraformaldehyde for 12 minutes at RT and washed with 1X PBS followed by permeabilization with 0.2% Triton-X-100 for 10 minutes at 4°C. Cells were blocked with 5% FBS and 0.05% Triton-X-100 and 0.2M glycine for 30 minutes at RT. Cells were incubated overnight with primary antibodies at 4°C followed by secondary antibody (Alexa flour 488 or Alexa flour 594) and DNA probing dye DAPI (Bio-Rad) for nuclear staining for 1 h at room temperature and mounted using Fluoromount G (#00-4958-02; Thermo Fisher Scientific) and images were acquired with confocal fluorescence microscopy. For live cell imaging, 1 x 10^6^ cells were seeded in 35 mm groove plates and required treatments were done for indicated time-points. Fresh medium was added after treatment with mitoSOX (#M36008, Invitrogen/Molecular Probes) and Hoechst 33342 (Sigma-Aldrich) for 30 minutes and analyzed by confocal fluorescence imaging. All Images were acquired using a Leica DMi8 inverted confocal laser scanning microscope (Leica-Microsystems) equipped with LED halogen light source for 405-, 488- and 594-nm laser sources and a 60× objective (oil immersion) using Leica Application Suite LAS software. Multiple z-stacks were captured using the software’s automatic scanning mode and maximum projected z-stacks as required. Images were acquired at room temperature in all experiments. Adobe Photoshop version 24.2.0 was use to convert images to 300 dpi format. Images were analyzed using IMAGEJ version 1.8.0_172 (64-bit) software.

### MTT assay

0.01 x 10^6^ cells were seeded in each well of 96 well cell culture plates. After 24 hours cells were preincubated with PJ34, EX527 before rotenone treatment for 24 hours. After treatment, cells were incubated with MTT reagent (#58945; Sisco Research Laboratories) at a specific concentration of 0.5mg/ml for 1 hour at 37°C. Upon appearance of coloration, the MTT solution was carefully aspirated and the formazan crystals were dissolved in DMSO. Quantification of formazan crystals in each well was done by measuring the absorbance in a microplate reader (BioTek, Synergy H1) at 540 nm. Each result of the experiment groups was normalized to the absorbance of the solvent control group for every assay.

### qRT-PCR analysis

Total RNA was extracted from cells following the manufacturer’s protocol for the PureZOL™ RNA Isolation Reagent (#7326880, Bio-Rad). cDNA was prepared by reverse transcription using iScript™ Reverse Transcription Supermix for RT-qPCR (#1708841, Bio-Rad) according to the manufacturer’s protocol. qRT-PCR was performed by iTaq Universal SYBR® Green Supermix (#1725124, Bio-Rad). The primers used were listed in **Table S2**, β2 microglobulin (B2M) were used as the internal control, and the relative expression of indicated genes was calculated using the 2^-ΔΔCT method.

### SIRT1 activity assay

For determining SIRT1 activity, the SIRT1 fluorometric assay kit (#BML-AK555; Enzo life sciences) was used according to manufacturer’s instructions. Briefly 1 x 10^6^ cells were lysed in 50μl of 0.5% NP-40 NETN (20mM Tris-HCl, pH 8.0, 100mM NaCl, 1 mM EDTA, and 0.5% Nonidet P-40) lysis buffer for 30 minutes in ice with periodic vortexing. Cell lysates were collected by centrifugation at 14000 rpm for 3 min at 4°C. Protein estimation was done by BCA assay kit and 25 μg of lysate was incubated at 37°C for 15 min. Volume for all the samples were adjusted to 35 μl with assay buffer (50 mM Tris-HCl, pH 8, 137 mM NaCl, 2.7 mM KCl, 1 mM MgCl2). 1mM DTT was added to each sample and further incubated for 15 min at 37°C. 15 μl of assay buffer with substrate, NAD and Trichostatin A were added and incubated for 1 hour at 37°C. Finally, 50 μl of 1X developer and nicotinamide solution added and incubated at 37°C for another 1 hour. Fluorescence was measured in white plate using a plate reader (Synergy H1, BioTek, Winooski, VT, USA) with excitation/emission of 360/460. Fluorescence readings were plotted for graphical representation of SIRT1 activity as Arbitrary Fluorescence Units (AFU).

### Cell cycle analysis

For analyzing cell cycle, SH-SY5Y cells were seeded in 6 well plate at a density of 0.3 x 10^5^ cells/well and respective treatments were done at 70% confluency. Following treatment, cells were collected by trypsinization and centrifuged at 200 g for 5 min. Further, the cells were resuspended in 300 μL 1X PBS and fixed in ethanol added dropwise while continuously vortexing at a low speed. Fixed cells were stored at 4 °C for 1 h, which was then centrifuged at 200 g for 5 min to discard the ethanol and washed thrice with 1X PBS at 300 g for 5 min. The cells were resuspended in 1X PBS to attain desired cell density necessary for the analysis and treated with RNase A (100μg/mL) for 1 h. Finally, for analysis, the samples were incubated in 40 μg/mL propidium iodide in dark for 30 mins and cell cycle distributions were analyzed using a BIO-RAD S3e cell sorter and was analyzed utilizing BIO-RAD FCS express flow cytometry analysis software.

### Reanalysis of publicly available ChIP-seq data

Peak calling and reanalysis were performed using *ChIP-Atlas* (https://chip-atlas.org) from Peak Browser, with a threshold of significance 50, with track type class set to TFs and track type to HMGB1. ChIP-seq peak profiles were visualized using the Integrative Genomics Viewer (IGV, version 2.19.1). Data were aligned to the hg38 reference genome. bigWig files representing signal intensity were imported, and peaks were displayed using a log scale with autoscaling enabled. For this purpose, GSM5234124 HUVEC cell line, GSM5234116 IMR90 cell line (34) and GSM2589815 HUVEC cell line (Zirkel et al.,2018) were used to identify the putative binding of HMGB1 on regulatory regions of DDR genes.

### Statistical analysis

All statistical analysis were carried out using GraphPad Prism version 8.0.2 for Windows (GraphPad Software, La Jolla, CA, USA). The results are presented as mean ± SEM of atleast three independent experiments or as indicated. Statistical significance was measured by one-way analysis of variance (ANOVA) with Tukey post-test or as indicated. Asterisks indicate level of significance (*P < 0.05, **P < 0.01 and ***P < 0.001).

### Declaration of AI in the writing process

During the preparation of this work the author(s) used ChatGPT (https://chatgpt.com/) in order to improve the English in certain cases and not for drawing any scientific conclusions.

## Supplementary information

**Fig. S1.**
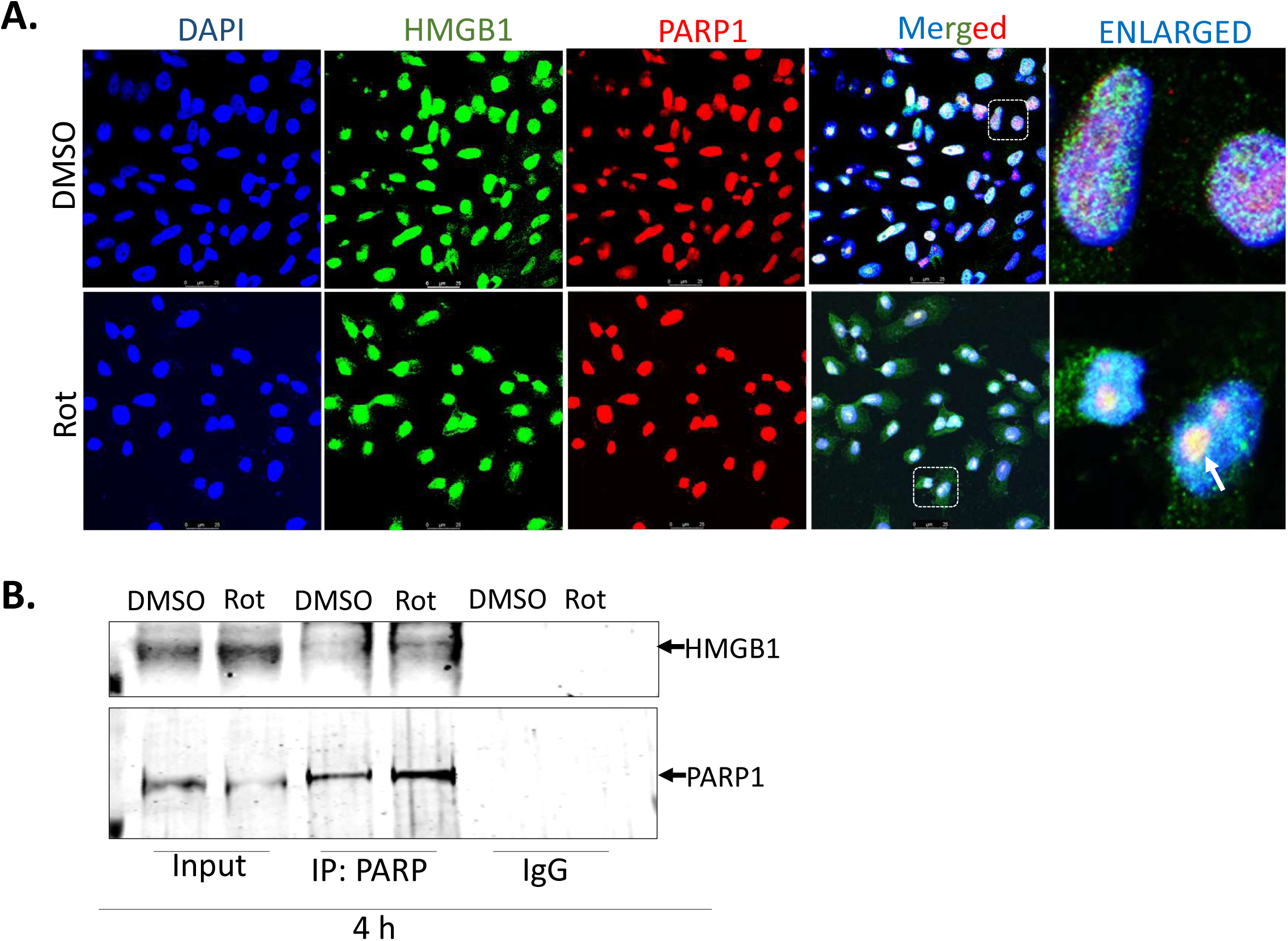
HMGB1 interacts with PARP1 following rotenone treatment. **(A)** Representative confocal images of co-localization of HMGB1 (green) with PARP1 (red) in cells treated with rotenone (5µM) or DMSO control for 4 h. DAPI (blue) indicates nuclear staining (scale bar=10 µm). **(B)** Co-immunoprecipitation analysis of HMGB1 and PARP1 in the whole cell extracts of cells following rotenone treatment for 4 h.

**Fig. S2.**
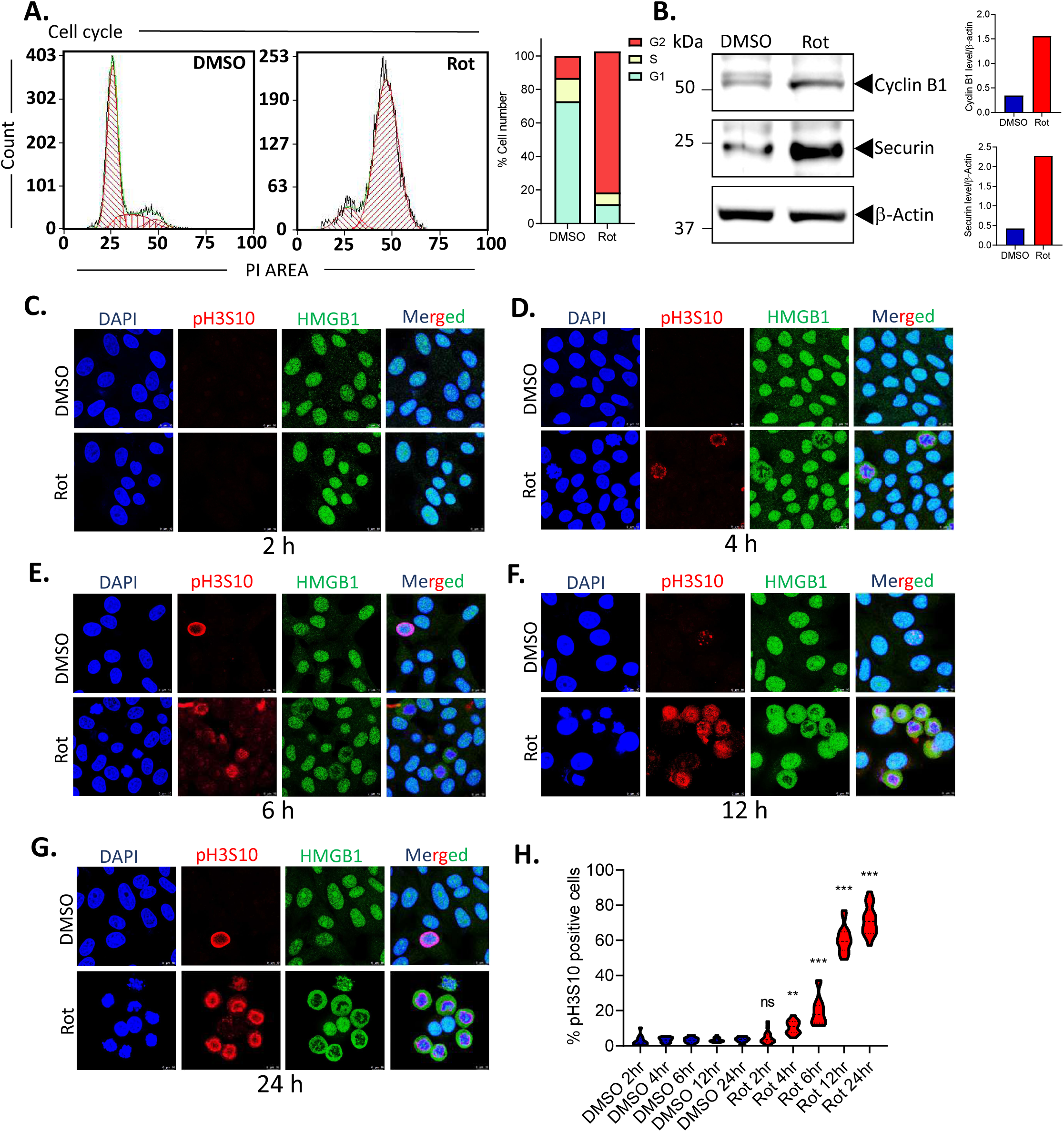
Rotenone induces G2/M arrest in SH-SY5Y cells. **(A)** Representative flow cytometry histograms depicting DNA content in cells stained with propidium iodide (PI). Distinct peaks correspond to cells in the G0/G1 phase (first peak), S phase (intermediate region), and G2/M phase (second peak). SH-SY5Y cells were treated with rotenone for 24 h and the data is compared with DMSO control. **(B)** Immunoblot analysis of Cyclin B1 and Securin protein levels in the whole-cell extracts of SH-SY5Y cells 24 h post-rotenone treatment. β-Actin was used as a loading control. **(C-G)** Representative confocal images of cells stained with pH3S10 (red), HMGB1 (green) and DAPI (blue) following rotenone treatment for 2 h (C), 4 h (D), 6 h (E), 12 h (F) and 24 h (G). (scale bar=25 µm). **(H)** Quantification of pH3S10 positive cells indicating cells arrested at G2/M for each treatment condition. Data are presented as mean ± SEM. Statistical significance was determined by one-way ANOVA with Tukey’s multiple comparison test (B) and was indicated by **p < 0.01; ***p < 0.001.

**Fig. S3.**
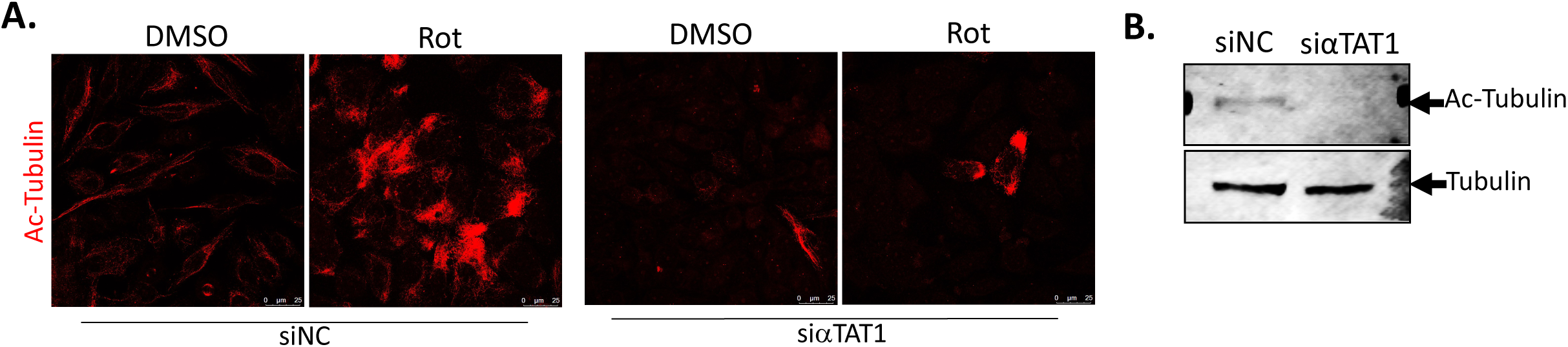
αTAT1 prevents tubulin acetylation following rotenone treatment. **(A)** Representative confocal images of cells stained with Ac-αTubulin (green) and DAPI (blue) in cells transfected with either siNC or siαTAT1 following rotenone treatment for 24 h (scale bar=25 µm). Results were compared with DMSO treated controls. **(B)** Immunoblot analysis of acetylated α-tubulin in the whole cells extracts of cells transfected with either siNC or siαTAT1. β-Actin was used as a loading control.

**Fig. S4.**
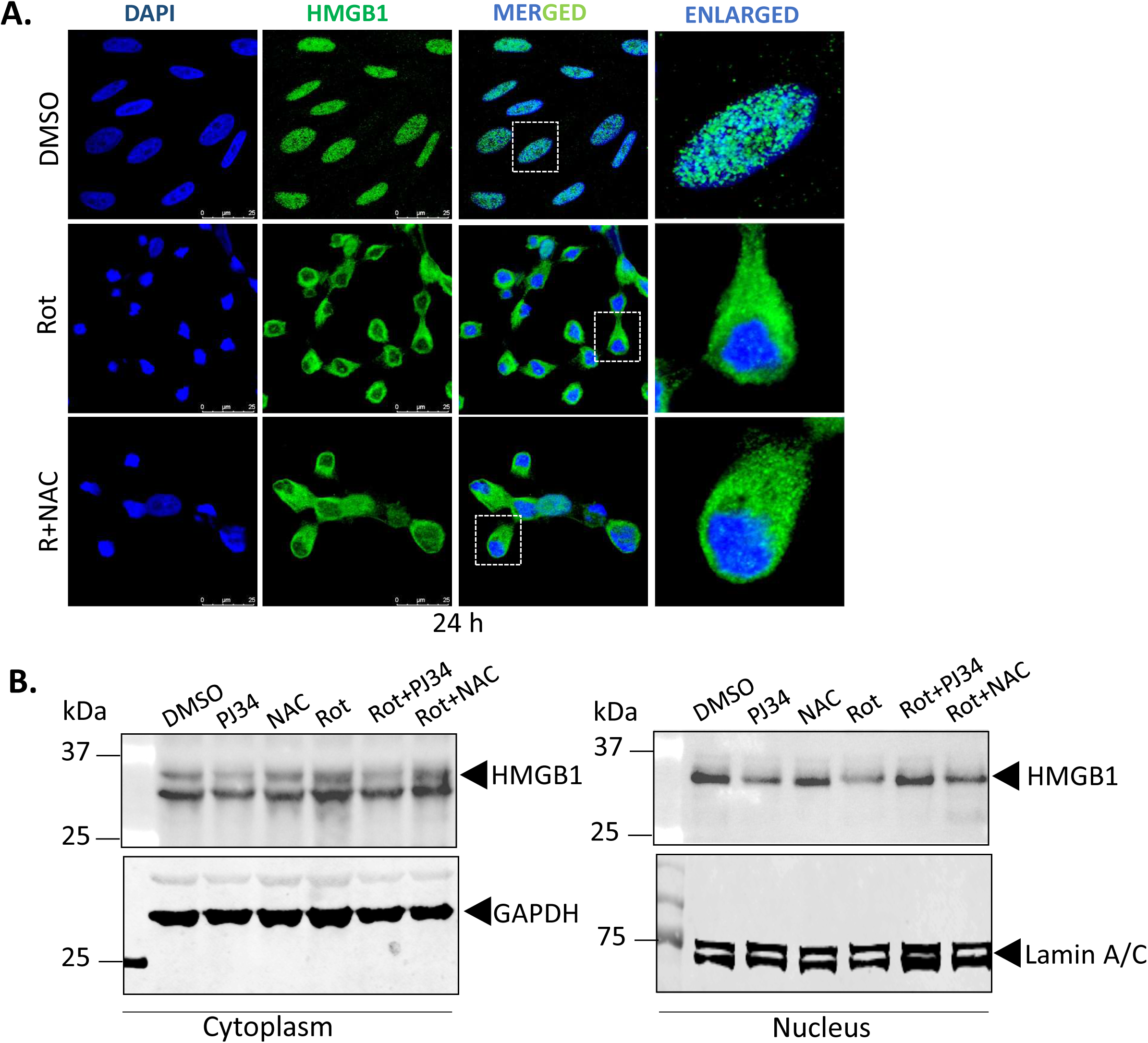
The anti-oxidant NAC did not prevent rotenone-induced HMGB1 nuclear exit. **(A)** Representative confocal images of HMGB1 (green) and DAPI (blue) in cells treated with rotenone for 24 h in the presence or absence of NAC. (scale bar=25µm). **(B)** Immunoblot analysis of HMGB1 protein levels in the nuclear and cytosolic fraction of cells treated with rotenone for 24 h in the presence or absence of the NAC. GAPDH was used as cytosolic loading control and Lamin A/C was used as a nuclear loading control.

**Table S1.**
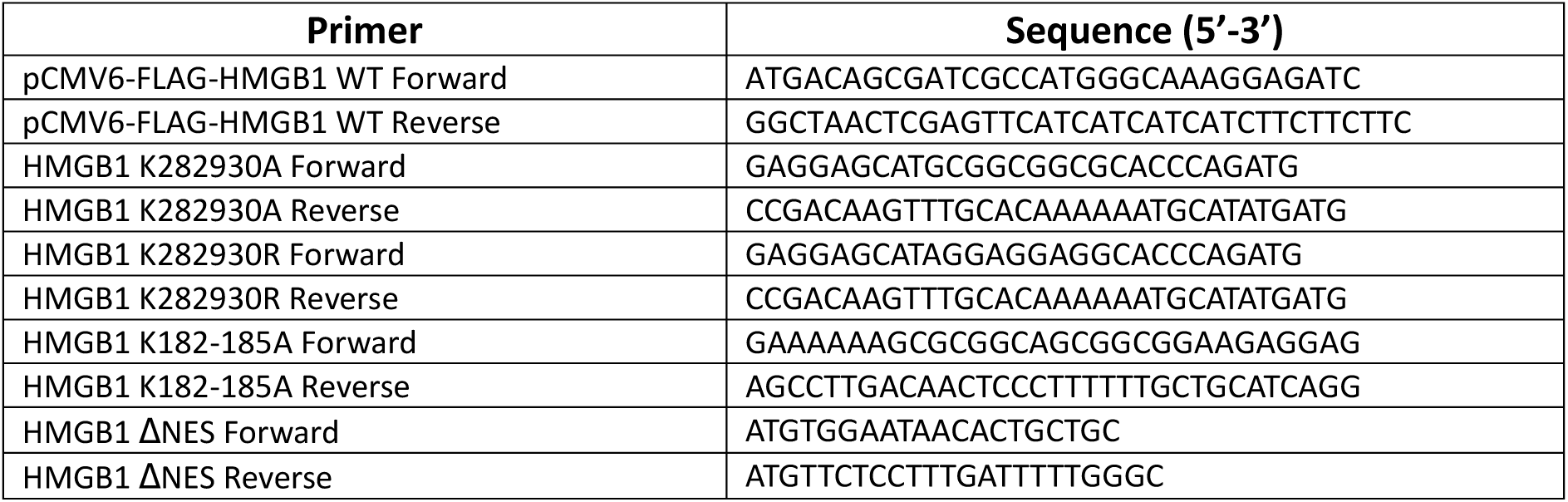
Primers used for cloning WT-HMGB1 and for site-directed mutagenesis.

**Table S2.**
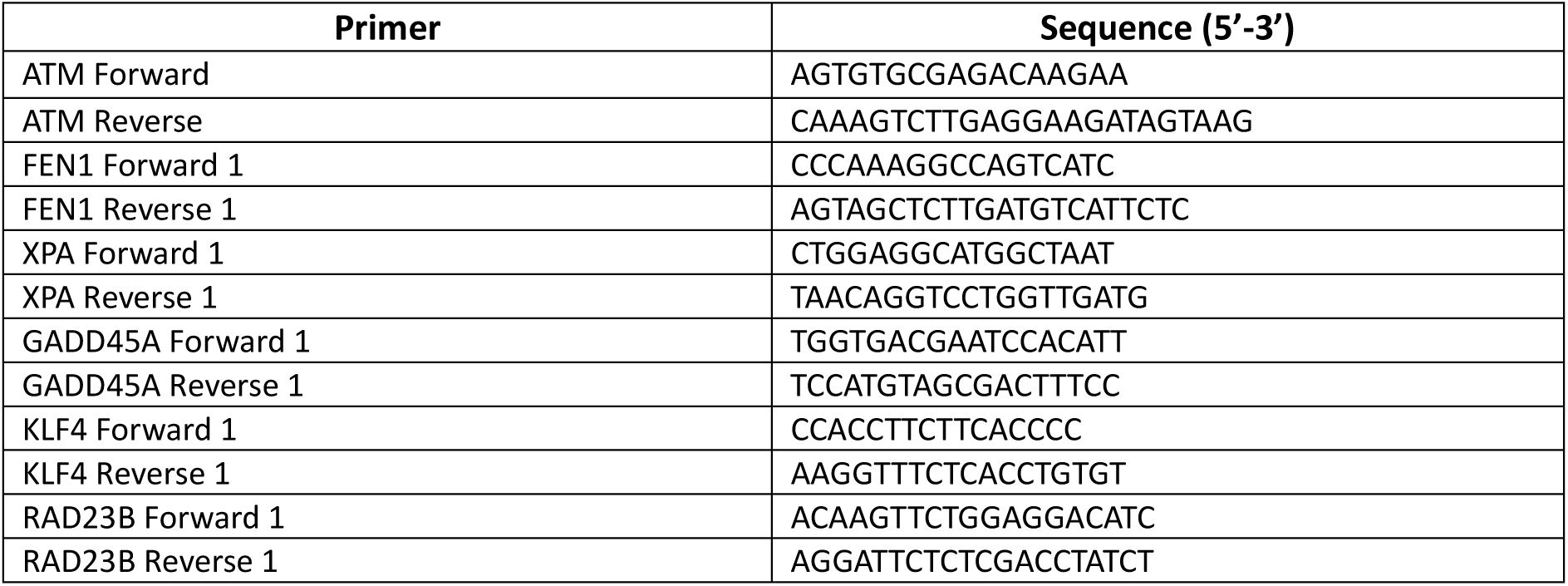
Primers use for qRT-PCR.

